# A practical framework RNMF for the potential mechanism of cancer progression with the analysis of genes cumulative contribution abundance

**DOI:** 10.1101/2021.08.12.456096

**Authors:** Zhenzhang Li, Haihua Liang, Shaoan Zhang, Wen Luo

## Abstract

Mutational signatures can reveal the mechanism of tumorigenesis. We developed the *RNMF* software for mutational signatures analysis, including a key model of cumulative contribution abundance (CCA) which was designed to highlight the association between genes and mutational signatures. Applied it to 1073 esophageal squamous cell carcinoma (ESCC) and found that APOBEC signatures (SBS2* and SBS13*) mediated the occurrence of *PIK3CA* E545k mutation. Furthermore, we found that age signature is strongly linked to the *TP53* R342* mutation. In addition, the CCA matrix image data of genes in the signatures New, SBS3* and SBS17b* were helpful for the preliminary evaluation of shortened survival outcome. In a word, *RNMF* can successfully achieve the correlation analysis of genes and mutational signatures, proving a strong theoretical basis for the study of tumor occurrence and development mechanism and clinical adjuvant medicine.

---

The cancer genomes contain many mutations, which are derived from exogenous and endogenous mutational processes that operate during the cell lineage [1]. These mutational processes are cumulative effects of DNA damage and repair processes, indicating unique patterns in tumorigenesis, namely mutational signatures [2–5]. Each mutational process from a tumor may involve some particular signatures that their biological combination processes could induce a large number of mutations [6–7]. Via studying the completeness of these mutations and identifying the digital genomic footprints that contribute to the mutation characteristics of tumors, we can not only reveal the potential mutational process information, understand the carcinogenic mechanism of tumor occurrence and development, but also provide biomarkers for early diagnosis, accurate cancer stratification and clinical response prediction, and realize individual treatment strategies [8–13]. Analysis of mutational signatures may reveal previously unknown mutation mechanisms and mysterious environmental exposure, such as herbal supplements containing aristolochic acid [13]. However, understanding of the pathogenic biological processes is still limited. Therefore, in order to systematically describe the mutational process leading to cancer, it is necessary to decipher the mutational signatures from the somatic mutation catalog by using mathematical statistical methods [14–23], the number of mutations that each feature in a single sample can be attributed to each feature, which annotate the probability of each mutation class in each tumor and the possibility of each feature producing.

Currently, the final referenced mutational signatures are archived in the catalogue of cancer of somatic mutations in cancer (COSMIC, https://cancer.sanger.ac.uk/cosmic/signatures). Most of them are common in various tumors, of which are specific to a certain type of tumor, of which are part of normal cell biology, and of which are related to abnormal exposure or tumor progression [24–31]. They may be attributed to known environmental exposure and mutation processes, such as tobacco smoke, ultraviolet radiation, the activity of the APOBEC series of cyclobutylaminases, and DNA mismatch repair defects or mutations in POLE. Besides, as known to us, the association between genes and mutational signatures was confirmed, and much focus were paid to study the role of hotspot mutations in the formation of mutational signatures, which provides a good research idea for the mechanism of tumorigenesis and development. However, at present, there are few tools systematically and in-depth mining the relationship between genes and mutational signatures [32, 33], which undoubtedly does not bring much convenience to the causal association between mutational signatures and genes.

Here, we first used the R language to design a simple and convenient package *RNMF*, which can directly start from the mutation data set to realize the correlation analysis of mutational signatures. Then, we pooled 1073 samples from Asian ESCC population, and then used *RNMF* to verify the practicability of this method framework. During the analysis, we highlighted interesting correlations through association analysis with driving mutations. Finally, deep learning method is used to explore the CCA matrix image data of gene under fixed features, and hierarchical learning of prognosis is done.

## Results

### Framework analysis of RNMF

Optimizing and improving the derivation of mutational signatures can not only rediscover some known features, but also produce new discoveries that were previously masked by technical and biological confusions. Here, we designed a simple and convenient process framework by R language (**Fig. 1**), and developed a novel R package called *RNMF* (https://github.com/zhenzhang-li/RNMF). The package *RNMF* can directly analyze the mutation data set which exist in the form of MAF or VCF format files, rapidly obtain the number of specific mutation types in each sample and then resolve the mutational signatures. This software can achieve 7 kinds of outstanding functions: (1) the mutation rate per M shown in LEGO graph; (2) rapidly extraction of mutational signatures; (3) accurately evaluating the contribution of each sample to the known mutation profile; (4) deeply studying on the similarity of mutational signatures and displaying the heat-map; (5) calculating the cumulative contribution abundance (CCA) matrix of genes; (6) studying the causal relationship between mutational signatures and genes; (7) studying on the relationship between hotspot mutation of driving factors and mutational signatures.

**Figure 1.**
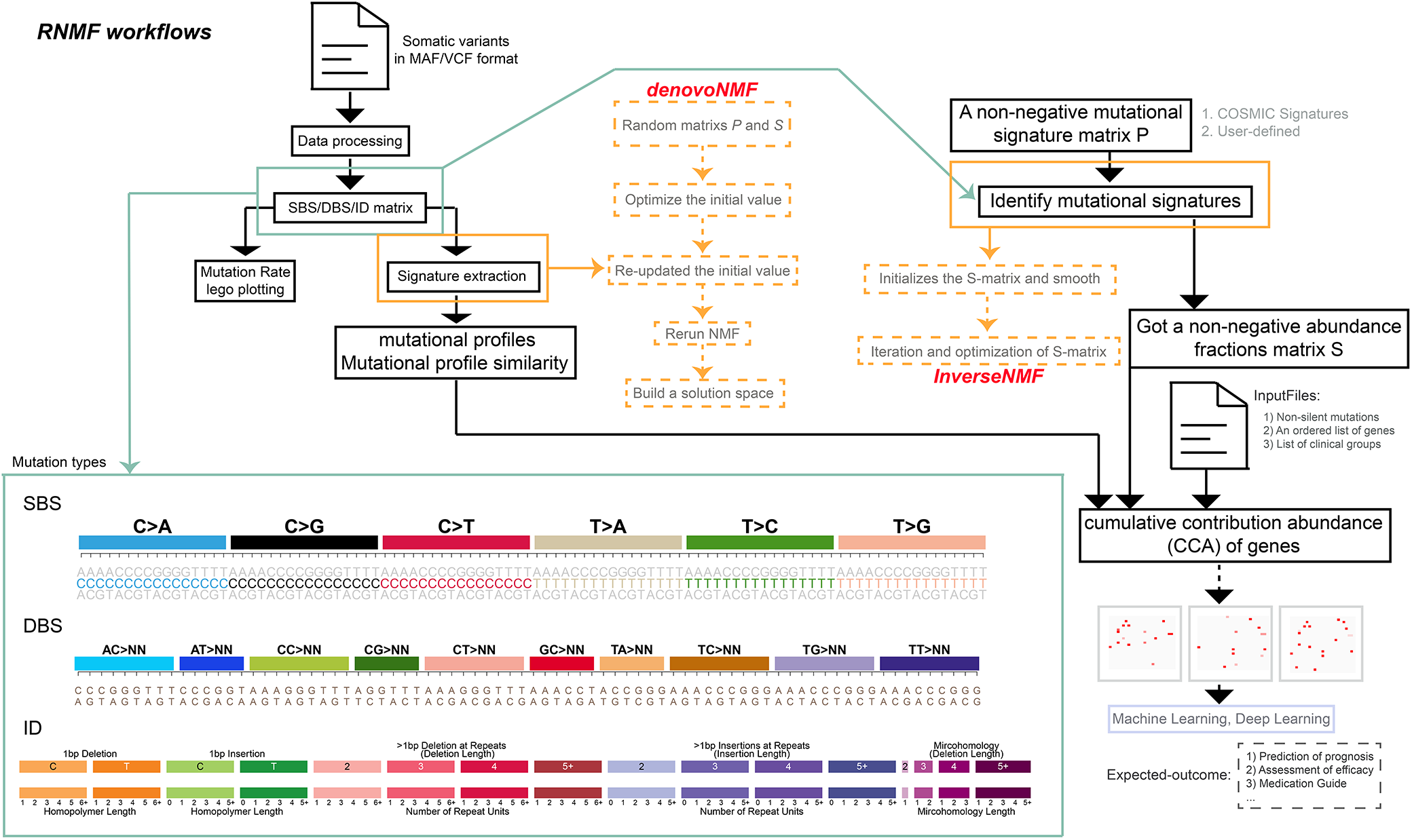
Overview of RNMF workflows. The automatic process can start from inputting mutation data files in MAF or VCF format to generate the necessary mutation type matrix, including SBS mutation types with 96 different contexts, 78 strand agnostic DBS mutation types, and a compilation of 83 different types of ID mutation types. Next, the process can use two program interfaces (*denovoNMF* and *InverseNMF*) for signature analysis to get the mutational profiles. Then, according to the clinical grouping, non silent mutation data and the list of genes to be studied, combined with P matrix and S matrix, the CCA matrix of genes was calculated. At the same time, the corresponding image data are generated according to the specified order, and finally the training and analysis are carried out by machine learning or deep learning methods, so as to obtain the expected results.

One outstanding function of *RNMF* is defined as *SigsInput,* which is suitable for large or most small datasets, providing analysis interfaces for different types of genomic DNA changes, such as single base substitutions (SBS), double base substitutions (DBS), small insertion and deletion mutation (ID). To prove high performance of our designed *RNMF*, the mutation data sets of 1073 ESCC samples from Asia (including 508 WGS data) are analyzed thoroughly. These mutation data were extracted from the appendix of published articles (**Table S1**), and were annotated by *Oncotator* [34]. As shown in **Fig. 1**, by extracting the mutational signatures with the designed *denovoNMF* function, the evaluation results exhibits the change of silhouette coefficient and error gradient. The criteria we chose are: 1) stable, without sudden decline or relatively large gradient of descent and large width of confidence interval; 2) small standard error term, and the gradient of standard error between adjacent classes tends to be gentle.

In the **Fig. 1**, an application interface “*InverseNMF*” can calculate the fraction of one sample which is contributed to each provided mutational signatures. Here, the threshold is set to 0.9. By our designed *RNMF*, we analyzed the contribution scores of these coding region mutations to each mutational signature which are gotten from the COSMIC Mutational Signatures (v2.0 - March 2015). The results are highly consistent with the analysis ones of the known “*MutationalPatterns*” [20] and “*deconstructSigs*” softwares [35], as shown in **Fig. S1a**. It is worth noting that operating speed of “*InverseNMF*” rivals to that of “*MutationalPatterns*” software, however, is 12 times faster than that of “*deconstructSigs*” software (**Fig. S1b**). To go insight into the inner link between mutational signatures and gene mutations, a series of powerful functions are designed according to the the CCA calculation model of gene, such as “*cumulativeCA*”, “*genePerMutSigs*”, “*samFisherSigs*”, “*samPerMutSigs*” and “*eachMutationCA*". The detailed analysis description will be provided in the example of our designed *RNMF*.

According to the previous reports [7, 24, 33], we define that if the CCA of a gene in a sample to a signature is less than 6%, the gene in the sample has little effect on the signature. Next, we combined non-silent mutations, sorted gene list and clinical grouping information to obtain gene mutation image sets of samples under different mutational signatures based on gene CCA matrix. Then, deep learning method was used to process these image information for hierarchical learning, which could obtain some biological cognition, so that we could deeply understand some useful information for adjuvant therapy, such as prognosis evaluation, efficacy evaluation, medication guidance and so on **(Fig. 1**).

In a word, as a versatile R package, *RNMF* can realize parallel operation that helps to study and evaluate the mutational processes during tumor development. Thus, molecular analysis can be performed based on extracted mutational signatures, further revealing the molecular mechanisms and optimizing the diagnosis and treatment decisions.

### Identifying mutational signatures via RNMF

The incidence and mortality of esophageal cancer have always been relatively high, as in China, for example, among which esophageal squamous cell carcinoma (ESCC) accounts for 90% of esophageal cancer [36]. A previous study has provided a large genome-wide sequencing cohort of Chinese ESCC population [37]. In order to demonstrate the role of *RNMF* in the extraction of mutational singatures, our designed *RNMF* is used to systematically analyze the mutation data set (single base mutations and INDEL mutations) of this cohort data. In the overall single base mutation pattern of ESCC, it is mainly C>T and C>G mutation, followed by C>A mutation, accounting for 34.9%, 18.81% and 15.94%, respectively (**Fig. S2**). Besides the insertion of a zero-length 1-bp T base homologous sequence, most other types of deficient incongruities are characterized by long (≥5) thymine mononucleotide repeats.

In this cohort, the *RNMF* successfully identified 12 single base substitutions signatures, named SBS1, SBS2, …, SBS12, respectively, which is compared with the COSMIC signatures (https://cancer.sanger.ac.uk/signatures; **Fig. 2, S3a, Table S2a-b**). Mutational Signatures description are represented in **Table 1**.

**Figure 2.**
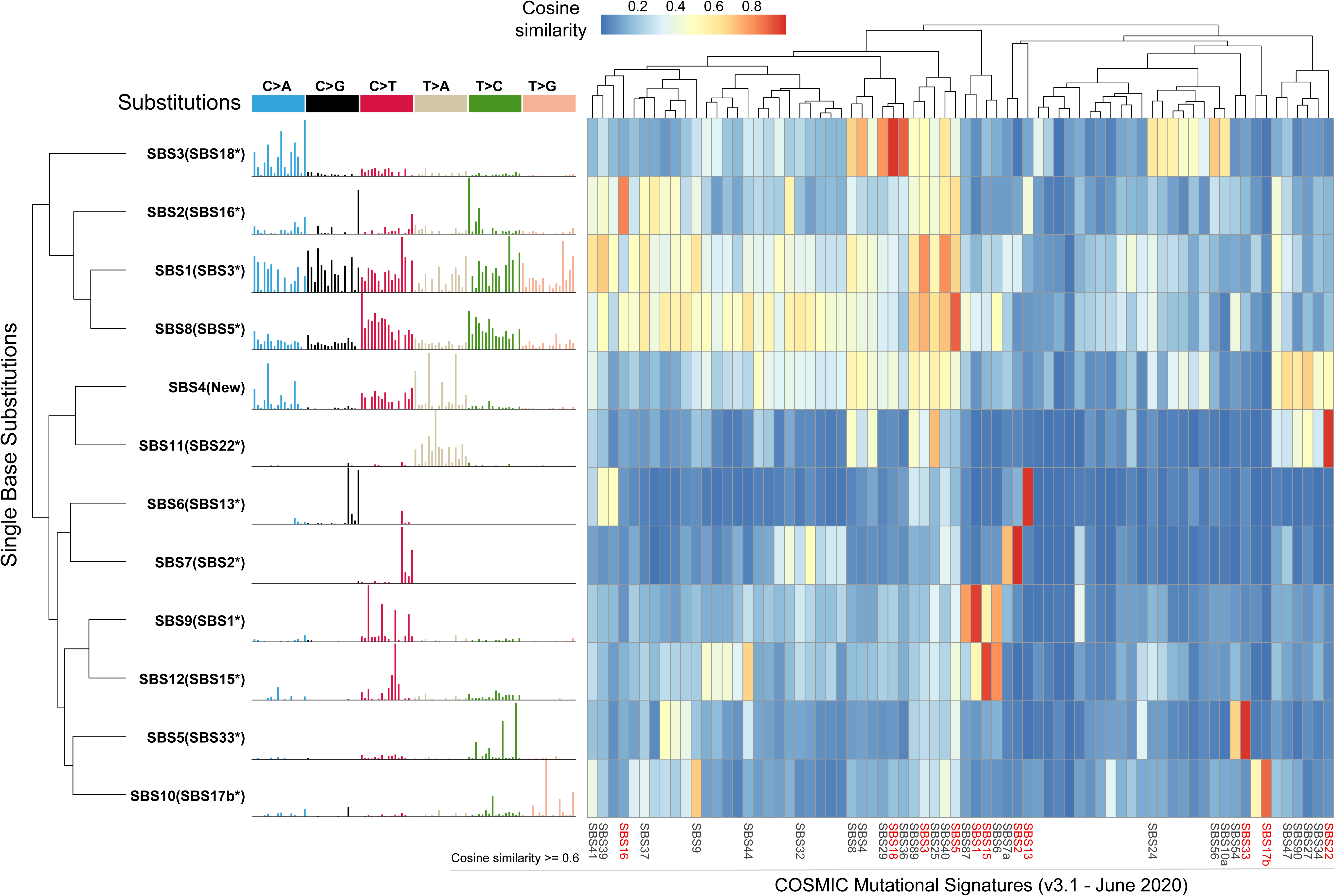
SBS signatures extracted from 508 Chinese ESCC patients. The left side of the picture shows the classifications of 96 mutation types. Each color is used to illustrate the positions of each mutation subtype on each plot. The right side of the picture shows heatmap of the cosine similarity between mutational signatures and COSMIC Mutational Signatures (v3.1 - June 2020). The shade of color corresponds to different cosine similarity scores. The SBS signatures with cosine similarity score no less than 0.6 are shown at the bottom of the figure, and the most similar one is highlighted in red.

**Table 1.** Mutational Signatures description.

| Original Name | Matching COSMIC | Cosine Similarity | Redefine Name | Comments |
| --- | --- | --- | --- | --- |
| **SBS Signatures** |  |  |  |  |
| SBS1 | SBS3* | 0.83 | **SBS3\*** | *BRCA1* and *BRCA2* mutations;<br>*BRCA1* promoter methylation;<br>homologous recombination deficiency |
| SBS2 | SBS16* | 0.84 | **SBS16\*** | Alcoholic consumption |
| SBS3 | SBS18* | 0.98 | **SBS18\*** | *CDH1* mutations [21]; *MUTYH* mutations |
| SBS5 | SBS33* | 0.98 | **SBS33\*** | Unknown |
| SBS6 | SBS13* | 0.98 | **SBS13\*** | ABOPEC |
| SBS7 | SBS2* | 1.00 | **SBS2\*** | ABOPEC |
| SBS8 | SBS5* | 0.90 | **SBS5\*** | *ERCC2* mutations; tobacco smoking |
| SBS9 | SBS1* | 0.96 | **SBS1\*** | Age |
| SBS10 | SBS17b* | 0.90 | **SBS17b\*** | Gastric acid reflux; fluorouracil (5FU)<br>chemotherapy treatment |
| SBS11 | SBS22* | 0.98 | **SBS22\*** | Aristolochic acid |
| SBS12 | SBS15* | 0.95 | **SBS15\*** | DNA mismatch repair deficiency |
| SBS4 | – | – | **New** | Unknown |
| **ID Signatures** |  |  |  |  |
| ID3 | ID6* | 0.94 | **ID6\*** | Homologous recombination-based repair |
| ID6 | ID2* | 0.99 | **ID2\*** | DNA mismatch repair deficiency |
| ID2 | ID14* | 0.88 | **ID14\*** | Unknown |
| ID1, ID4, ID5,<br>ID7-ID9 | – | – | **New1-New6** | Unknown |

Note: “–” represents there is no match feature or the similarity is generally very low.

To verify the accuracy of the above results, the known *SigProfilerExtractor* software [23] is also used to analyze SBS signatures, and extract 13 features (**Fig. S3b, S4**). By further making a similarity comparison, we found that whether using V2 or V3 version of COSMIC signature, it shows strong consistency (**Fig. 2, Fig. S5a-b**). Moveover, the results were basically consistent between *RNMF* and *SigProfilerExtractor* (**Fig. S5c**). Thus, these results enough show the validity of the *RNMF*. In addition, we also extract ID signature. Nine ID signatures were prominent (**Fig. S3c, Fig. 3, Table S2c-d**), and three of them have been previously reported, including two with known mutational processes [23]. Mutational Signatures description are represented in **Table 1**.

**Figure 3.**
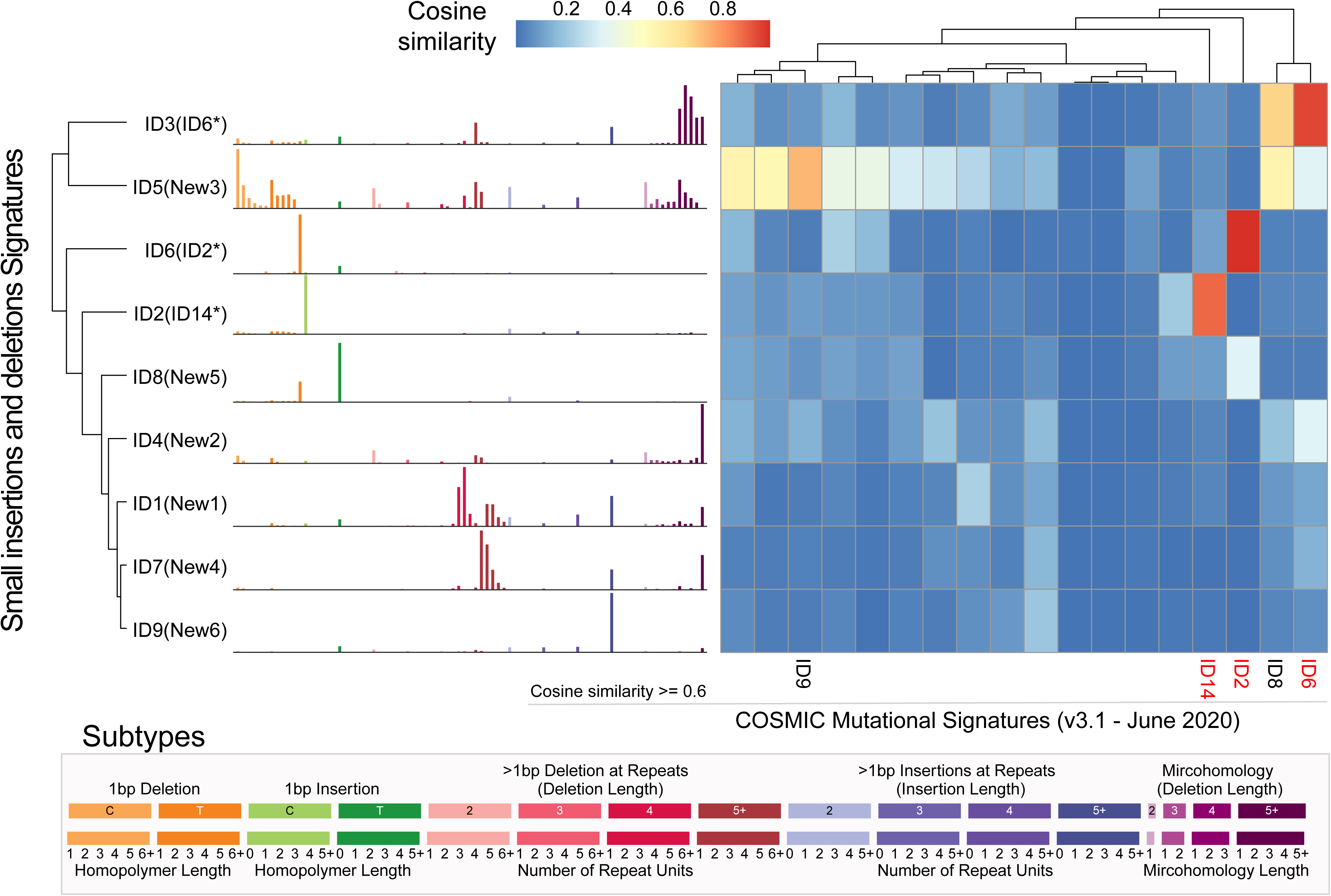
ID signatures extracted from 508 Chinese ESCC patients. The left side of the picture shows the classifications of 83 mutation types. Each color is used to illustrate the positions of each mutation subtype on each plot. The right side of the picture shows heatmap of the cosine similarity between mutational signatures and COSMIC Mutational Signatures (v3.1 - June 2020). The shade of color corresponds to different cosine similarity scores. The ID signatures with cosine similarity score no less than 0.6 are shown at the bottom of the figure, and the most similar one is highlighted in red. At the bottom of the figure, the specific information of 83 mutation types is given, and the colors correspond to the columns in the left image one by one.

### Estimation of sample contribution under known mutational signatures by RNMF

In order to verify the practicability of the functions in our framework, the contribution of samples was analyzed in the known mutational signatures. We collected a total of 115,130 somatic mutations from exon region of 1073 ESCC samples, including 16 of them were hyper-mutated with mutation count more than 500 (**Fig. 4a, Table S3**). Comparing with 508-WGS cohort, we found that the overall proportion of C>T mutations increased in the exon region, among which *[C>T]G context changed greatly (**Fig. S6a**). However, the single base mutations in the exon region were mainly C>T (48.19%) and C>G (18.2%), followed by C>A (13.8%), indicating that the mutation pattern of exon region was similar to that of whole genome.

**Figure 4.**
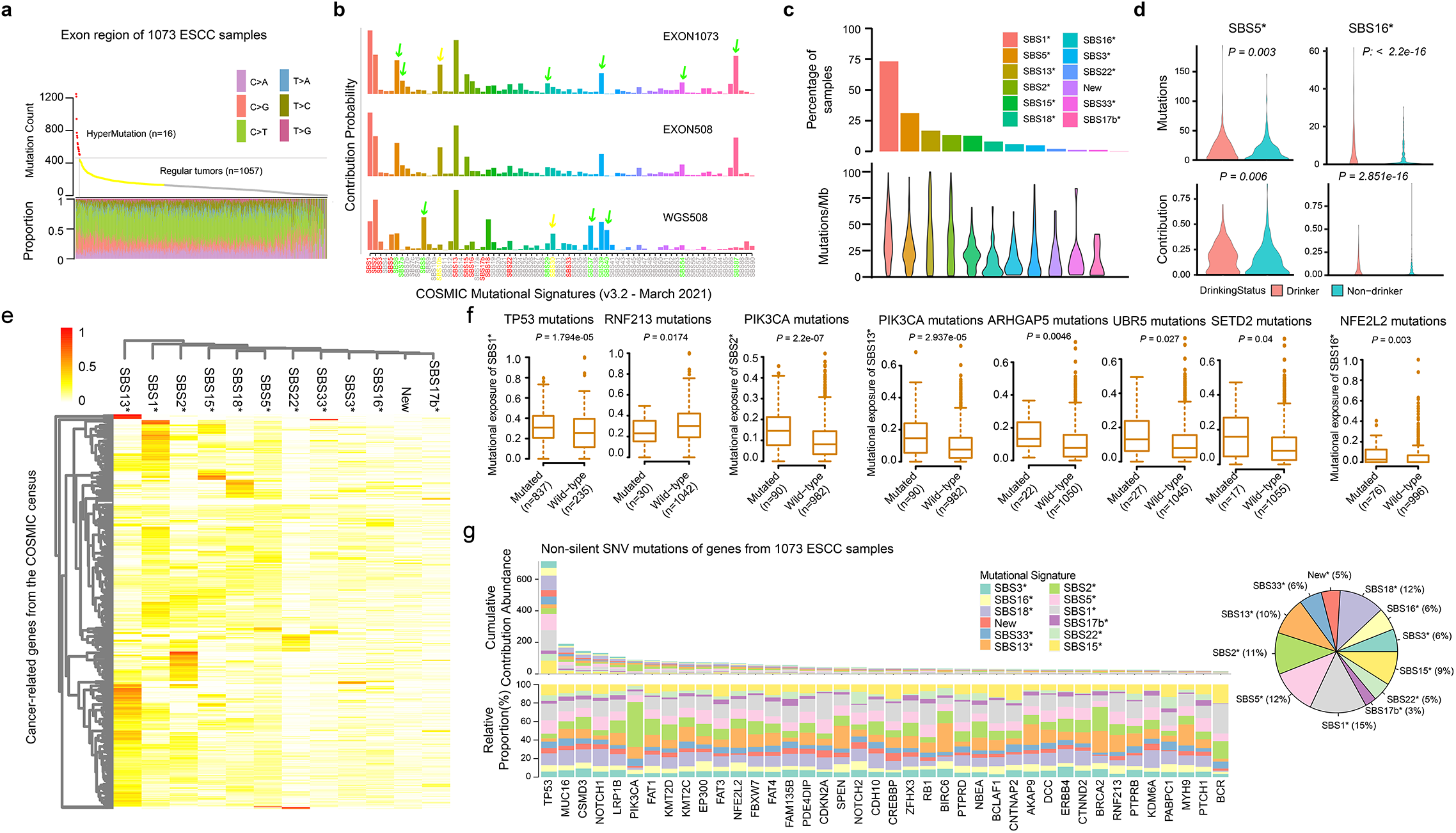
Mutational signatures reconstruction from 1073 ESCC cases. (a) The top figure of the graph shows the statistics of hypermutation. Using Ckmeans.1d.dp to cluster the number of mutations of 1073 ESCC samples, 16 hypermutated samples (red dots) and 1057 regular tumors (yellow and gray dots) are found. The bottom figure of the graph shows the proportion of six mutation types of 1073 ESCC samples with point mutation, and the X-axis represents the sample, Each sample has a single column, and each color represents a mutation type. (b) Based on the background mutation contribution probability of COSMIC Mutational Signatures (v3.2 - March 2021), each color represents a mutational signature, the length of each column represents the contribution ratio of mutation to the signature, the red mark represents the signature most similar to the 12 mutational signature, and the green arrow and green font indicate that this signature is very similar to the 12 mutational signature. Yellow means that the similarity between the signature and 12 mutational signatures is very low. (c) Percentage of ESCC tumors in each signature was displayed (top) and mutation rate for each signature in the relevant samples (bottom). If the contribution of a sample assigned to one signature is not less than 20%, we would consider that this signature is present in the sample. (d) The association between mutational activity of SBS signatures (SBS5* and SBS16*) and alcohol consumption. The violin compared the difference between drinking and non drinking groups from the mutation count and the contribution of samples to the signatures, and the significant p value was statistically analyzed by Student’s T-test with two-sided. (e) The heatmap shows the distribution of CCA of cancer-related genes from the COSMIC census, and the depth of color represents the degree of correlation. (f) Box plot showing that the SBS signatures were associated with cancer-related genes mutations (including SNV and indel), where n represents the number of samples. Statistical significance was tested by rank sum test with two sided. (g) The contribution of non-silent mutations in the coding regions of 40 cancer-related genes is statistically analyzed. Each color represents a class of mutation feature map, and the pie chart shows the proportion of each feature. In the figure, the top figure shows the total CCA of each gene in 1073 samples, while the bottom figure shows the proportion of each mutational signature, and a column represents a gene. Gene selection rules: the number of non-silent mutations is more than 30 and belongs to cancer-related genes from the COSMIC census.

To better explore and analyze the potential features of exon regions, we used COSMIC Mutational Signatures (v3.2 - March 2021 and v2.0 - March 2015) as the background to obtain the number of contribution mutations of samples in each signature. We analyzed three single base mutation datasets: 508-WGS cohort (WGS508), exon region of 508-WGS cohort (EXON508) and exon region of 1072 samples (EXON1073). For the 12 feature maps extracting from WGS data, the trends of WGS508, EXON508 and EXON1073 are basically the same under the same background (**Fig. 4b, S6b**). However, the results of WGS508 revealed that the signature, which accounted for a large proportion, may not exist in the list of 12 decomposed signatures, and these signatures either have high similarity or low similarity related to those 12 signatures. What’s more, the results of WGS508 showed that compared with the results of different versions of COSMIC Mutational Signatures, we found that with the increase of features background analysis, high similar or low similarity signatures with high proportion will be produced, indicating that adding background features may introduce some unimportant or similar features to share the load weight **(Fig. 4b, S6b**). Meanwhile, under the same background, the results of EXON508 and EXON1073 showed that the number of samples play little role in the proportion of signatures (**Fig. 4b, S6b**). In addition, previous reports [24, 37] showed that each kind of cancer has its own important characteristics. Of which the number or type of signatures is usually different. These different main signatures play a leading role in the occurrence and development of different tumor types.

In this paper, by combining the results of our analysis and the reported results (**Fig. S4**), we strongly believe that 12 stable SBS signatures, which is extracted from the 508-WGS cohort, should be the outstanding features of ESCC, which are considered as the leading mutation patterns in the occurrence and development of this tumor. Therefore, these 12 features were selected as background walls for the analysis of ESCC tumors (**Table S3**). We found that the signatures SBS1*, SBS2*, SBS5*, SBS13* and SBS15* were effective in most samples containing more somatic mutations, which are considered as ubiquitous signatures (**Fig. 4c, S6c**). By contrast, signatures SBS3*, SBS16*, SBS17b*, SBS18*, SBS22*, New, and SBS33* were sporadic signatures which exsit in rare samples (≤8% of cases). Interestingly, clinical association analysis revealed that compared with non-drinking patients, drinking patients contributed significantly more mutations to SBS5* and SBS16*, and their contribution also increased significantly, suggesting that these two SBS signatures may be related to alcohol consumption (**Fig. 4d**).

### CCA analysis reveals potential prognostic features and mechanisms through RNMF

In this work, a total of 717 cancer-related genes are screened from the COSMIC census (https://cancer.sanger.ac.uk/census, **Table S5**) and their CCAs were calculated via *RNMF* software with a given model function named *cumulativeCA*. Through this program algorithm interface, CCA of a gene on a signature and CCA of a gene on a signature in a sample can be obtained. We found that these cancer-related genes are more enriched in ubiquitous signatures, especially signatures such as SBS1*, SBS2*, and SBS13* (**Fig. 4e**). There are differences in CCA levels of genes under different mutational signatures, which indicates that genes have their preference for mutational signatures. At the same time, survival analysis found that CCA of several genes was associated with prognosis (**Fig.S7a, Table S6**). In addition, a multivariate Cox model was confirmed that some of them were still significant (**Fig. S7b**), such as *ARHGAP5*, *SETD2*, *RNF213*, *CDKN2A*, *NOTCH1*, *NFE2L2* and so on. Analysis of mutation characteristics showed that *TP53* mutation significantly increased exposure to SBS1*, but conversely, the contribution to SBS1* was significantly reduced in samples carrying *RNF213* mutation (**Fig. 4f, S8a**). Furthermore, among the *TP53* mutant samples, the contribution of the RNF213 mutant samples to SBS1* was significantly reduced(**Fig. S8a**). Similarly, we found a significant increase in the contribution of APOBEC signatures (SBS2*, SBS13*) in ESCC samples with *PIK3CA* mutations, and the other three genes (*ARHGAP5*, *SETD2* and *UBR5*) were also associated with APOBEC signature (SBS13*) (**Fig. 4f, S8a**). It is noted that *NFE2L2* mutation was related to SBS16*. These above results indicate that there is a potential mechanism between gene mutations and mutational signatures in the process of tumorigenesis and development. Thus, we studied 40 representative genes from 717 cancer-related genes, which contained about 3.6% of the total number of non-silent SNV mutations. Most of these mutations preferred the characteristic SBS1* (15%), SBS2* (11%), SBS5* (12%), SBS13* (10%), and SBS18* (12%), but were less distributed in SBS17b* (3%) (**Fig. 4g**). We found that 66.5% of 1073 ESCC samples had non-silent SNV of *TP53*, resulting in a highest level of CCA of *TP53* gene, and the proportion of three SBS signatures (SBS1*, SBS5* and SBS18*) was higher, followed by SBS15*. Obviously, different genes have different ratios for different mutated traits. It attracts our attention that *PIK3CA* gene is obviously in favor of APOBEC signatures (SBS2*), which accounts for more than 50% (**Fig. 4g**). Hence, all findings above suggest that preference of cancer-related genes for mutational signatures can be defined by CCA, which can further expose some potential prognostic features or mechanisms.

### CCA analysis exposing PIK3CA helical mutation E545K are strongly mediated by APOBEC

According to the reports [38–40], *PIK3CA* is a typical proto-oncogene that typically harbors some hotspot mutations in tumors and is enriched in APOBEC characteristics in a variety of cancer types, especially these two most-common and well-studied hotspots: E542K (c.1624G>A) and E545K (c.1633G>A) in the helical domain. In ESCC, the cohort results of previous studies implicitly implicated APOBEC activity as a key driver of *PIK3CA* mutagenesis [40, 41]. In this cohort, we are committed to further study the potential mechanism between *PIK3CA* biological mutations and APOBEC signatures via CCA model.

By the CCA enrichment analysis, it is found that tumors with non-silent mutations in *PIK3CA* had increased activity of the signature SBS2* (ESCC1073-EXON: 89 tumors with non-silent *PIK3CA* mutations and a median increase CCA of 0.74 per sample; q=0, *P*=0; Regular tumors of ESCC1073-EXON: 88 tumors with non-silent *PIK3CA* mutations and a median increase CCA of 0.74 per sample; q=0, *P*=0; WGS508-EXON: 38 tumors with non-silent *PIK3CA* mutations and a median increase CCA of 0.741 per sample; q=0, *P*=0; **Fig. 5a, S8b**). Analogously, those tumors with non-silent mutations in *PIK3CA* also had increased activity of the signature SBS13* (ESCC1073-EXON: 89 tumors with non-silent *PIK3CA* mutations and a median increase CCA of 0.136 per sample; q=0, *P*=0; Regular tumors of ESCC1073-EXON: 88 tumors with non-silent *PIK3CA* mutations and a median increase CCA of 0.1359 per sample; q=0, *P*=0; WGS508-EXON: 38 tumors with non-silent *PIK3CA* mutations and a median increase CCA of 0.133 per sample; q=0.0008, *P*=0.0001; **Fig. 5b, S8c**). In order to further prove these connections, we performed mutational signature enrichment analyses and gained the same results (**Fig. S8b, Fig. S8c**), providing the strongest statistical evidence for the relationship between *PIK3CA* mutation and APOBEC signatures activity in ESCC. Together, these results further strongly suggest that, although APOBEC signatures activity are present in all tumors, somatic *PIK3CA* mutations are associated with a significant increase in APOBEC signatures activity.

**Figure 5.**
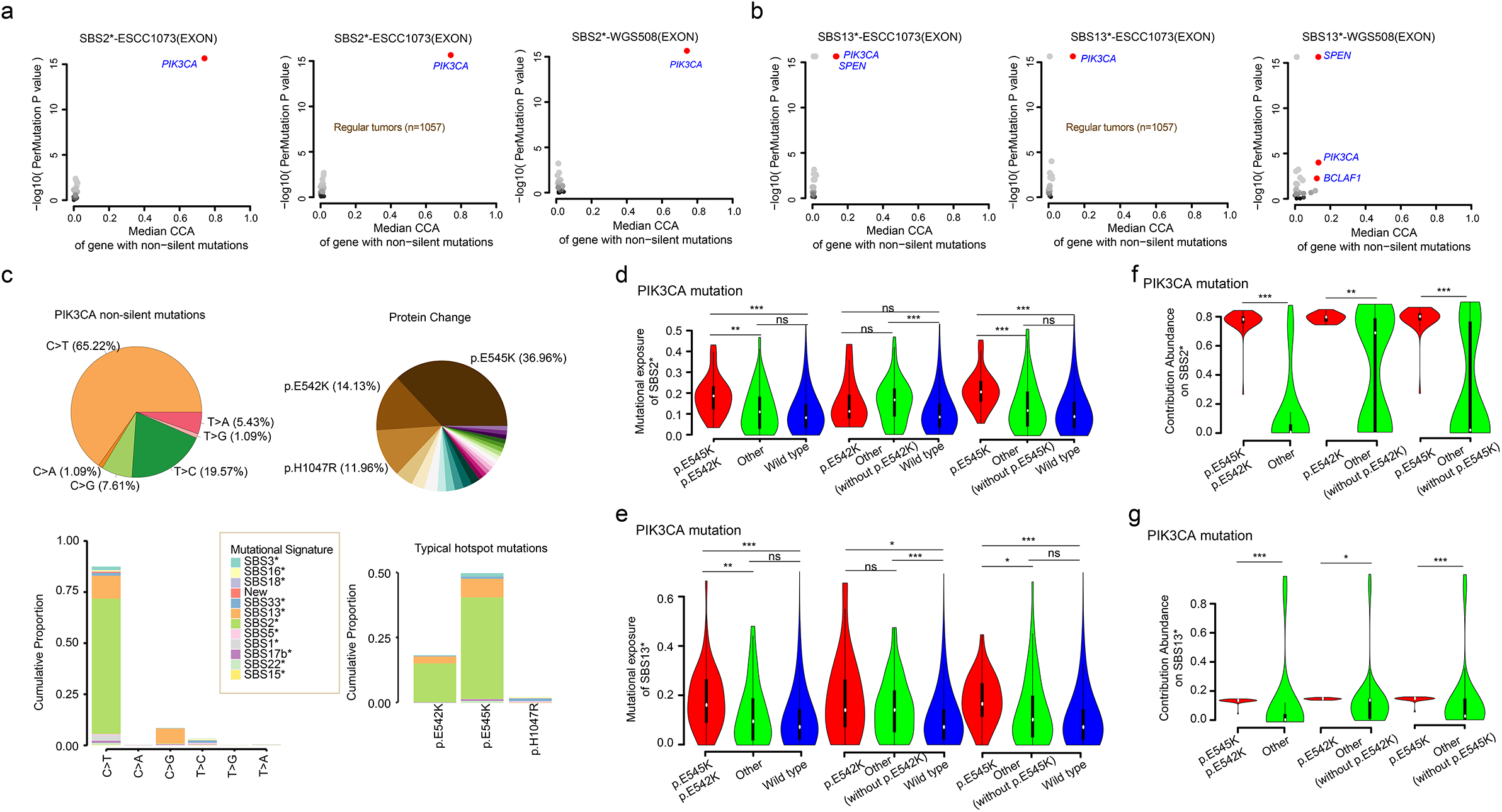
CCA enrichment analysis identifies an association between somatic PIK3CA mutations and activity of SBS2* or SNS13* in ESCC. (a-b) Here, we use two datasets: exon regions of 1073 and 508 ESCC cases. First, the median CCA of each gene in the current signature is calculated, and then the contribution importance of each gene is calculated by PERMUTATION test to study the association between gene and signature. The regular patterns in the figure represent the samples with non hypermutated. For genes mutated in >5% of samples, the CCA of genes attributed to SBS2* or SBS13* was compared in tumors with wild-type versus mutated copies of the gene. Genes with FDR q < 0.1 are highlighted in red. (c) Mutation trend and hotspot analysis of *PIK3CA* non-silent mutations: the pie chart on the top left shows the proportion of six mutation types, and the pie chart on the top right shows the proportion of coding protein, with the name of the protein that accounts for a large proportion. Each color in the figure below represents a mutational signature, and the statistical proportion of the contribution of six mutation types to each mutation signature is on the left, a column represents a mutation type; The figure on the right shows the contribution abundance of classical hotspot mutant protein to each signature, and a column represents a classical mutant protein. (d-e) The violin diagram shows the difference of contribution to SBS2* and SBS13* between the samples with *PIK3CA* hotspot mutation and other types of samples, while (f-g) shows the difference of contribution abundance between the samples with *PIK3CA* non-silent mutations (E545K and E542K) and other samples. The Wilcoxon rank sum test with two-sided is used here, * represents (0.01 < = P < 0.05), * * (0.001 < = P < 0.01), ***represents (P < 0.001) and ns represents (P > = 0.05).

We also dissect the mutation spectrum of *PIK3CA* non-silent mutations, revealing the underlying mechanism of during mutation processing. In our series, 65.22% of *PIK3CA* non-silent mutations were C>T substitution, and those mutations were frequent presenters that mostly contributed to APOBEC signatures (SBS2* and SBS13*) with highest percentage (**Fig. 5c**). Simultaneously, we investigated the *PIK3CA* helical (E545K: 36.96%; E542K: 14.13%) and kinase (H1047R: 11.96%) hotspot mutations, and found that only the helical mutations had a high cumulative proportion for APOBEC signatures (SBS2* and SBS13*)(**Fig. 5c**). Then, We related the *PIK3CA* helical mutations to each APOBEC signatures, and observed a significant increase for mutational exposure of APOBEC signatures in samples harboring helical domain mutations (**Fig. 5d-e**). Significantly, tumors carrying a hotspot mutation E545K significantly hold a high degree of contribution fraction of SBS2*, yet hotspot mutation E542K can not bring significant benefits to SBS2*, as well as the other mutations (**Fig. 5d**), implying that only mutation E545K can affect the benefit of the overall mutation data of the sample on SBS2* compared with other mutations. Similarly, we found that although the E542K mutation significantly increased the benefits of SBS13*, the significant intensity of the increase was not as high as that of E545K (Median: 0.146 vs. 0.172) (**Fig. 5e**), indicating that the E545K mutation in *PIK3CA* can accelerate the increase of SBS13* activity. Furthermore, from the perspective of gene itself, the CCA of *PIK3CA* genetic hotspot mutations for APOBEC signatures was significantly higher than that of other mutations (**Fig. 5f-g**). However, compared with E542K mutation, the effect of E545K mutation is more significant, indicating E545K among *PIK3CA* mutation is more closely associated with APOBEC signatures. In a word, *PIK3CA* helical mutation E545K contributes more significantly to APOBEC signatures, suggesting that they are strongly mediated by APOBEC.

### CCA analysis displaying the relationship between age signature and TP53 typical hotspot mutations

In previous study, the results of *TP53* mutations on mutational signatures indicates that driver mutations of *TP53* mutations are associated with specific mutation processes in human cancers, such as colon, skin, bladder, lung, and liver cancers [42, 43]. They only mentioned that the most frequent *TP53* mutations were associated with the most commonly observed age signature which featured by C>T transitions at CpG dinucleotides. It is worth considering that there is no detailed report on the association between *TP53* typical hotspot mutations and age signature in the related studies of ESCC, including the previous large cohort analysis of ESCC [37, 44]. Here, we analyzed the association between *TP53* mutations and age signature confirming that age signature was associated with *TP53* mutations via CCA enrichment analysis (Regular tumors of ESCC1073-EXON: 715 tumors with non-silent *TP53* mutations and a median increase CCA of 0.0823 per sample; q = 0, *P* = 0; ESCC1073-EXON: 728 tumors with non-silent *TP53* mutations and a median increase CCA of 0.085 per sample; q = 0, *P* = 0; **Fig. S8d**). Moreover, mutational signature enrichment analyses also revealed the strong relationship between *TP53* mutation and age signature activity (**Fig. S8d**). It’s worth noting that *TP53* was mainly enriched with C>T substitutions (44.51%) with a large proportion of them were preferentially contributed to age signatures (SBS1*)(**Fig. 6a**). We screened the six kinds of hotspot mutations with the highest risk rate (R342* : 4.07%; R213*: 3.37%; R282W : 2.95%; R175H: 2.81%; R273H: 2.53%; R248Q: 2.11%) and analyzed their association with mutational signatures. We found that except R175H, the other five hotspots preferred the age signatures (SBS1*) (**Fig. 6a**), which indicated that there was a potential mechanism between these hotspots and age signatures (SBS1*). Compared with other mutations, tumors harboring at least one of these hotspots will significantly increase its contribution to age signature (SBS1*) (**Fig. 6b**). Actually, although R282W can improve the contribution of the sample to age signature (SBS1*) (R282W vs. Other Mutation vs. Wild-type: median increase of 0.367 vs. 0.306 vs. 0.276), only hotspot mutation R342* can significantly affect the benefit of the whole mutation data of the sample to age signature (SBS1*) (R342* vs. Other Mutation vs. Wild-type: median increase of 0.388 vs. 0.305 vs. 0.276; **Fig. 6b**), indicating that R342* mutation is the primary factor to increase the activity of age signature (SBS1*). However, from the perspective of gene mutation itself, the CCA of *TP53* typical hotspot mutations for age signatures was significantly higher than that of other mutations (**Fig. 6c**), which indicated that except for R342* and R282W mutations, the other four hot spot mutations could not significantly improve the overall mutation contribution level of the sample, but they are still involved in the specific mutational process. In addition, under age signature (SBS1*), tumors with CCA of *TP53* no less than 0.06 were significantly associated with poor prognoses (*P*=0.44, **Fig.S7a**). Notably, CCA of *TP53* in tumors carrying *TP53* mutation R248Q were more than 0.06, and those cases were associated with deceased survival outcomes (**Fig. S8e**). Multivariate cox model shows that *TP53* hotspot mutation R248Q are independent prognosticators for poor survival in ESCC (**Fig. S8f**). Finally, we also found that *TP53* small INDEL mutations were related to New3 (ID signature; **Fig. S8g**).

**Figure 6.**
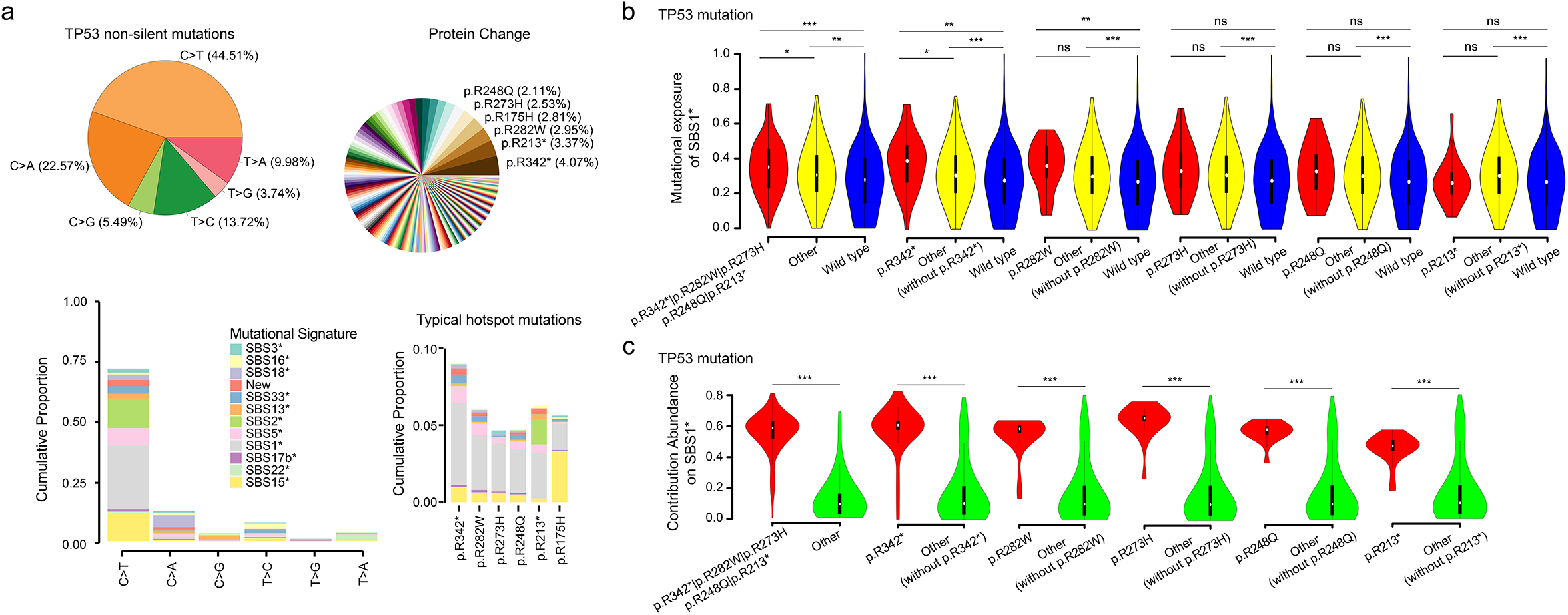
CCA enrichment analysis identifies an association between somatic TP53 mutations and activity of SBS1* in ESCC. (a) Mutation trend and hotspot analysis of *TP53* non-silent mutations: the pie chart on the top left shows the proportion of six mutation types, and the pie chart on the top right shows the proportion of coding protein, with the name of the protein that accounts for a large proportion. Each color in the figure below represents a mutation feature, and the statistical proportion of the contribution of six mutation types to each mutation signature is on the left, a column represents a mutation type; The figure on the right shows the contribution abundance of classical hotspot mutant protein to each signature, and a column represents a classical mutant protein. (b) The violin diagram shows the difference of contribution to SBS1* between the samples with *TP53* hotspot mutation and other types of samples, while (c) shows the difference of contribution abundance between samples with *TP53* hotspot mutations and other samples. The Wilcoxon rank sum test with two-sided is used here, * represents (0.01 < = P < 0.05), * * (0.001 < = P < 0.01), ***represents (P < 0.001) and ns represents (P > = 0.05).

### Image analysis based on CCA matrix of genes

To obtain some favorable statistical information, and even get some prognosis evaluation or medication guidance, which is helpful for clinical treatment, we designed a way to transform CCA of gene in each mutational signature into intuitive image information. The detailed operations are as follows: 1) we can sequence these genes through some potential relationships, such as similarity, pathway, or clinical association; 2) according to these CCA matrix results, some standard visual impact images are generated; 3) we can apply these images to deep learning model, combined with clinical information for analysis and mining.

In this work, we use “hclust” clustering to get the gene sequence based on the CCA matrix of genes. From our mutation data set, we will screen out the list of all non-silent mutation genes in the existing mutation set and do intersection processing with 717 genes. The intersecting gene set is the final data information transformed into image. We organize the data set into a *N*×*N* matrix (where 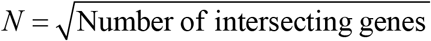). In this paper, we arrange the genes from top to bottom with the aim of obtaining the effect picture of gene mutation. To study the effect of CCA changes in each mutational signature on the prognosis, the deep learning framework PyTorch [45] was performed to train some publicly available models, such as ResNet [46] (ResNet50, ResNet101), DenseNet [47, 48] (DenseNet-121, DenseNet-161), Inception-V4 [49] and MoblieNet-V2 [50], V3 [51]. First of all, we uniformly limit the pixels of the picture to 500*500. Because G1 individuals are living and their follow-up time is less than 3 years, we will use G2-3-4 groups for the next analysis and mining, as shown in **Table S7a**. Limited by clinical information, and in order to better evaluate the prognosis, we especially compared G2 and G4 groups. For choosing an ideal model, we randomly select a feature as a template for training, and take the high average accuracy as the judgment basis for model selection. Here we choose one SBS signature (New) as an example, as shown in **Table S7b**. Four of them have higher accuracy, and they are Resnet50 (68.571%), DenseNet121 (71.429%), MoblieNetV2 (68.571%), and InceptionV4 (68.571%). In order to test the stability of our model, the above four models were trained for 10 times. In the training process, the model parameter random seed was fixed, other parameters are the same. Finally, through the analysis, we found that DenseNet-121 model has a higher average accuracy (**Table S7b; Fig. S9a**). Consequently, we chose DenseNet-121 as a model to analyze all signatures, the full schematic representation as shown in **Fig. S9b**. In the process, a stochastic gradient descent method [52] was used with an initial learning rate of 0.01, weight decay of 10-4 and momentum of 0.7 in the process of training. Next, dropout, data augmentation and L2-regularization were applied to prevent overfitting. The above parameter sets were properly tuned for DenseNet-121 model. Then, for testing the stability of data and finding the global optimal solution, the model DenseNet-121 was trained 10 times for each sub-feature data of G2-G4. Random seed was set free in the training process. That some less accuracy than others may be a local optimal solution, because stochastic gradient descent method was used as the optimizer in the training process. We found that the accuracy was comparatively stable to each sub-feature and relatively higher in mutational SBS3*, New and SBS17b* (**Fig. S9c**), suggesting that the beneficial feasibility of this conversion method of the CCA matrix image data of gene. Simultaneously, the results are given the best accuracy of G2-G4 is the sub-feature SBS17b* (77.500%), followed by SBS3* (72.500%) and New (71.429%). Furthermore, the probability distribution over the above 3 sub-features of G3 group in the G2-G4 was tested. Interestingly, the distribution of G3 group is more likely to fall on G2 group (**Fig. S9d**). So further the model DenseNet-121 was also trained 10 times for each sub-feature data of G2G3-G4 (**Fig. S9c**). The results show that the best accuracy of G2G3-G4 is the sub-feature SBS3* (76.190%), followed by New (72.973%), SBS16* (71.111%) and SBS17b* (70.732%), as shown in **Table S7c**. This illustrate that the G3 group addition has slight effect on the classification results. Additionally, the mutational signatures such as SBS3*, New and SBS17b* still have a high degree of explanation for G2-G4 and G2G3-G4 (**Fig. 7**). Finally, we found that the survival group or the samples with a follow-up time of no less than 3 years had a higher contribution to SBS3* and SBS17b*, respectively (**Fig. S9e**).

**Figure 7.**
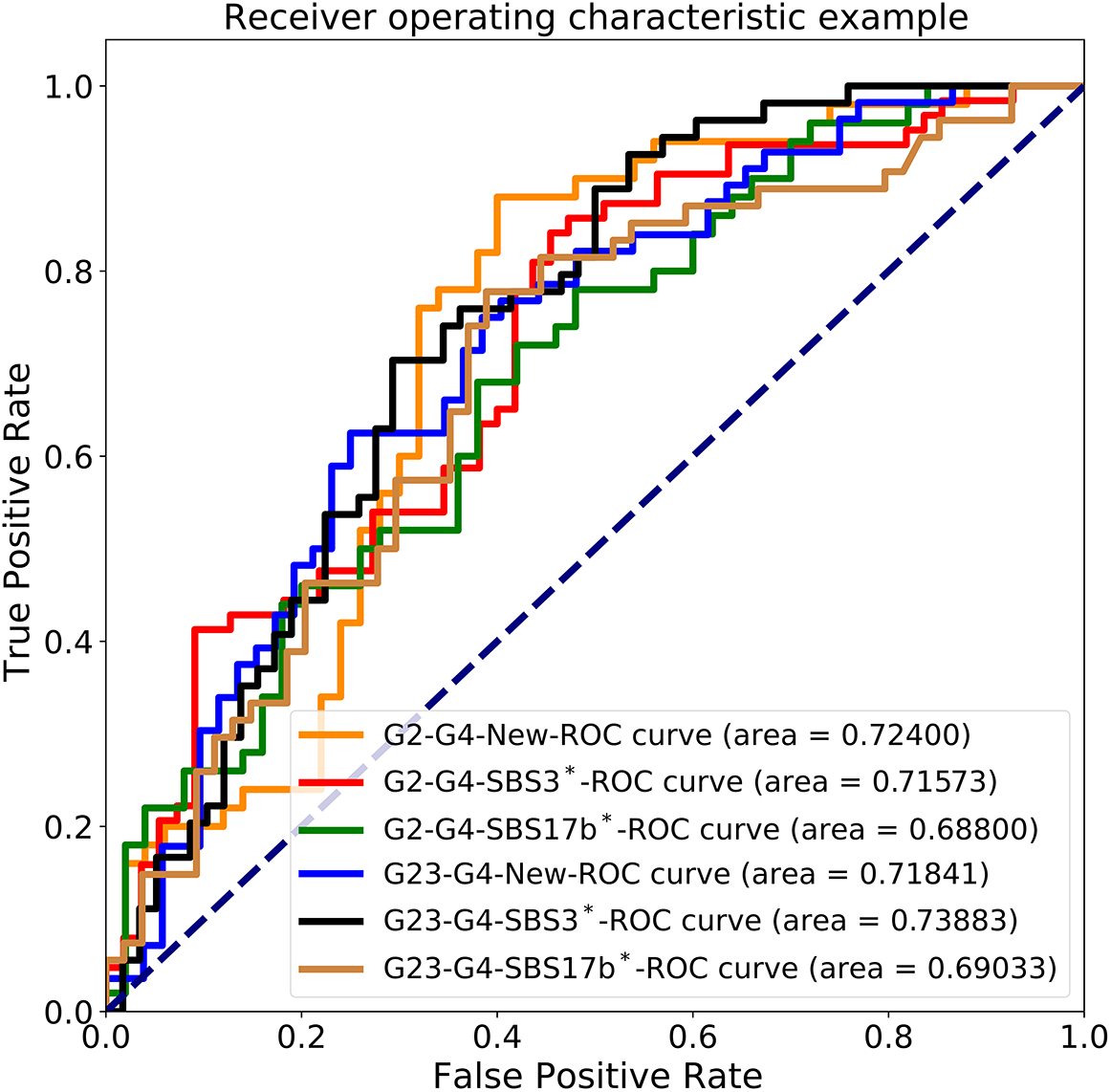
Application of CCA matrix of genes in prognosis for ESCC. The sensitivity-specificity curve of set classifier at the test sample sets for Urgent versus Non-urgent binary classification.

## Discussion

Here, we provide an integrated mutational signature analysis framework with a CCA model of genes, achieve a meta-analysis of 1073 ESCC samples, and verify the practicability and application value of our framework. Via this framework, we obtained known and uncovered previously undescribed signatures (including 12 SBS signatures and 9 ID signatures) from 508 WGS tumors of 1073 ESCC cases. And further identified and highlighted an association between *PIK3CA* helical mutation E545K and activity of APOBEC signatures. Similarly, we also reported that age signature and the hotspot mutation R342* of *TP53*, and *TP53* (R248Q) is a poor predictor for ESCC. In addition, the CCA matrix image data of genes under mutational signatures New, SBS3* and SBS17b* were calculated. This is helpful for the preliminary evaluation of short-term prognosis.

In addition to feature extraction and sample contribution analysis, we can also assign graph variation features to each sample by the designed framework, or even each gene, and then CCA of gene under a certain signature can be also obtained. Yet, compared with the previously published software [23], our framework spends more time on a de novo extraction of signature analysis. The reason is that we design a correction process “Re-updated the initial value” and a solution space process “build a solution space”. Hence, this is a weakness in the framework that need to be optimized in the future. However, our framework provides a new idea for understanding the panorama of tumor occurrence and development, and help scientific researchers to study the mechanisms of tumor progression. It is of great application value to study the characteristics and statistical distribution of one gene under a certain signature by assigning the signature. In this study, based on previous reports [23, 24, 37] and statistical evaluation (**Fig. 4b, S6b**), we confirmed the background of expected mutational signatures of ESCC. Concurrently, 717 cancer-related genes from the COSMIC census (https://cancer.sanger.ac.uk/census, **Table S5**) were selected as the base to calculate CCA. Due to the inconsistent sequencing background of the data, we uniformly analyzed the data of coding regions, and calculated the contribution of each sample to these signatures for subsequent analysis. However, this analysis will have some limitations. In order to avoid the impact of this limitation, we are committed to explore the ubiquitous signatures such as APOBEC signatures and age signatures, and discuss those frequently mutated genes that present in ESCC, such as *TP53* and *PIK3CA*.

Previous reports have revealed that mutations in the helix domain and kinase domain of PIK3CA cause activation through different mechanisms, and the mutation process may be related to driving mutations in a variety of cancers [53]. In ESCC studies, it was also mentioned that *PIK3CA* mutation was associated with the APOBEC signatures [40, 43]. Furthermore, we found that there was a close relationship between APOBEC signatures and *PIK3CA* mutations in the meta-analysis of 1073 ESCC tumors, especially *PIK3CA* mutation E545K. In conclusion, APOBEC mediated driver mutations, especially the known hotspot mutation E545K of *PIK3CA*, suggests that the activity of APOBEC is the main source of *PIK3CA* mutation in ESCC and an important factor in tumor development.

Next, we analyzed the association of *TP53* hotspot mutations and mutational signature. Notably, the most frequent *TP53* mutations found in ESCC were associated with the most commonly observed mutational signature, age signature, which reflects the natural degradation of 5-methylcyto sine to thymine [54]. In particular, the mutation R342* of *TP53* can affect the mutation process of tumor occurrence and development, resulting in a significant increase contribution of the sample to age signature (**Fig. 6b**). This led us to put forward the hypothesis that the mutation R342* of *TP53* in ESCC which is the primary factor to increase the activity of age signature. In many tumor types, driver mutations of *TP53* appear to be strongly associated with multiple signatures, and their probably arises due to the selection of loss-of-function (LOF) and dominant-negative (DN) alleles which are generated by specific mutational processes [41, 42]. In our analysis, *TP53* mutations R213* was also shown to be an independent prognostic factor.

In addition to the CCA matrix image, deep convolutional neural network denseNet-121 was used to analyze and the CCA matrix image data of gene in SBS3* and SBS17b*, which can be preliminarily distinguished the shortened survival outcome (follow-up time no less than three years). This finding indicates that the results of this method serves as one of the criteria to evaluate the prognosis of three-year survival. Combined with AI technology, our designed scheme directly provides a new way to explore from the single gene relationship research to the multi gene association analysis. In general, the number of individuals studied in this paper is relatively small, which is one of the shortcomings of model learning. We hope to further achieve the useful information in the era of big data. More excellent learning model is also one of the improvement ideas to obtain accurate results, which needs further research in the future. At the same time, complete clinical information, including treatment methods, medication information, extended follow-up time and so on, is of great clinical significance for the further exploration of this idea. We are reasonably optimistic that in the future, CCA matrix of genes can be used to evaluate the prognosis, metastasis risk, recurrence risk, and even provide medication guidance and suggestions for individuals.

Overall, it is indispensable to understand and explore the mechanism of tumorigenesis and development by studying the relationship between genes and mutational signatures. The potential application of CCA of genes needs to be further studied and explored, such as giving some specific gene lists, forming image pictures, and perhaps evaluating prognosis and guiding medication through deep learning.

## Method

### Genomic data collection

All somatic mutations were initially collected from the supplementary data of six previous studies (See **Table S1**) comprising 1073 esophageal squamous cell carcinoma (ESCC) cases, including 508 genome-wide data and 564 exon sequencing data.

### R package link and parsing description

In the analysis process of this software, somatic variants can be imported from a Variant Call Format (VCF) file or a Mutation Annotation Format (MAF) file. Then it relies on the Bioconductor library, such as BS.genome.Hsapiens.UCSC.hg19 or BS.genome.Hsapiens.UCSC.hg38, to acquire an information matrix of mutation types (SBS, DBS and ID, etc). Subsequently, the program extracts mutational signatures according to the generated data, and finally obtains the result files. The R package is publicly available at https://github.com/zhenzhang-li/RNMF. The detailed document file also provides some examples of commands usage. In addition, scripts for running the package will also be provided in the R package.

### Optimal mutational signature extraction framework

Based on the model definition in previous reports [55, 56], we can obtain a classical formula *V* = *PS* + *E*, which is used to extract the mutational signatures in human cancers. In this equation, *V* refers to the observation matrix with size *M*×*N*, of which *M* represents the observed characteristics, and *N* is the number of samples. Supposing the number of mutational signatures is *K*, then we can estimate a non-negative mutational signature matrix *P* with size *M*×*K* and a non-negative abundance fractions matrix *S* with size *K*×*N*. Simultaneously, error matrix *E* that refers to nonsystematic errors and sampling noise is calculated during the processing.

For an observed mutational catalogs *V*, these *K* mutational signatures could be extracted by *denovoNMF* (**Fig. 1**) as following:

#### Step 1 (Capture random matrices)

We randomly generate matrices *P*(*P*≥0) and *S*(*S*≥0). Ideally, for a mutational signature, its components are basically fixed, so the sum of its standardized components is equal to 1; for one sample, the total of normalized abundance fractions for each mutational signature should be infinitely close to 1. Hence, here we require that 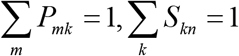

#### Step 2 (Optimize the initial value)

We apply resampling to obtain a new matrix 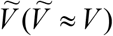 by using the Dirichlet distribution, and straighten matrices *P* and *S* by columns and then merge them into a solution vector **x**. An optimized objective function 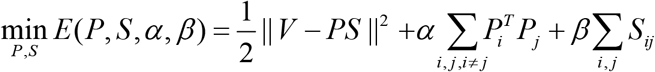 which previously reported [55] is used to find the best solution. After smoothing 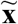, the updated matrices *P* and *S* are restored according to the straightening rules. Finally, we generate a new initial mutational signature matrix *P* and a new initial abundance fractions matrix *S* through 20000 iterations via referring to the previous implementation process [15].

#### Step 3 (Re-updated the initial value)

Perform Steps 1 and 2 for *I* (*I* ≥ 5) iterations. Their errors generated by iterations are calculated by formula *E* = || *V* − *PS* ||^2^, and the results of the 5 items with the smallest error are selected. Then we apply the k-means [57] algorithm to the set of matrices *P* and *S* to cluster the data into *K* clusters, respectively. Subsequently, class-center 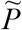 and 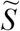 are obtained. Similar to Step 2, we straighten them to calculate the optimal solution space, and finally gain the excellent initial value.

#### Step 4 (Rerun NMF)

In this step, we still use the multiplicative update rules to generate the final matrix. The iterative model is as follows:

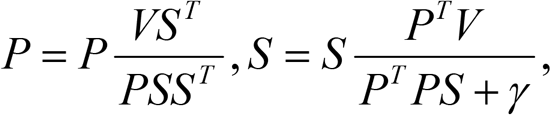

Where *γ* is a parameter to control the accelerated convergence. Iterate until P and *S* convergence or until the maximum number of 100,000 iterations is reached.

#### Step 5 (Build a solution space)

Repeat the process of steps 1 to 5 with 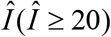 times to generate a solution space for all the value of *K* (*K* ∈ *N* ^+^). Then suitable selection in this solution space using the silhouette coefficient measure and error gradient.

After a series of analysis process, the final mutational profiles of each *K* clusters can be acquired. Follow the previous experience [20], we use the cosine similarity to determine the similarity between two mutational signature A and B.

### Deriving the contribution of defined mutational signatures

At the same time, we also developed a reversible method named *InverseNMF* (**Fig. 1**), which can identify mutational signatures within a small dataset or a single tumor sample. Previous reports [20, 33] have confirmed the importance of such applications and provided another analytical strategy for tumor characteristic map studies.

In this step, the *P* matrix is a user-defined matrix (such as the signature matrix provided by COSMIC), and then a feasible sample contribution matrix *S* can be estimated iteratively by observing matrix *V*. The iterative model is as follows:

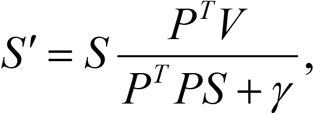

where *γ* is a parameter to control the accelerated convergence. Iterate until *S* convergence or until the maximum number of 100,000,000 iterations is reached.

### Mutational signature operative in ESCC

We applied our framework to extract mutational signatures from 508 WGS samples of 1073 ESCC tumors. At the same time, the previously reported tool “*SigProfilerExtractor*” [23] was used to analyze the data. Finally, the similarity between the two results was analyzed (**Fig. S5**). Our framework optimizes the non negative matrix factorization (NMF) algorithm and takes the change of silhouette coefficient and error gradient as the evaluation index of feature number selection (**Fig. S3a,c**).

In order to analyze and explore the potential features of exons of 1073 ESCC samples, We used COSMIC Mutational Signatures (v3.2 - March 2021 and v2.0 - March 2015) and mutational signals extracted from 508 WGS data as the background to obtain the number of mutations in each mutational signatures. Based on the results of three single base mutation data sets (508-WGS cohort renames as WGS508, exon region of 508-WGS cohort renames as EXON508, and exon region of 1072 samples renames as EXON1073), the eigenvalues of mutational signatures for subsequent analysis of ESCC were determined.

### Cumulative contribution abundance (CCA) of genes

As described earlier [7], since each mutational process takes into account the mutation category and the generation of mutations in tumor is attributed to each corresponding process, we define that the influence of mutations of category *m* in tumor *n* during the mutational process can be expressed as:

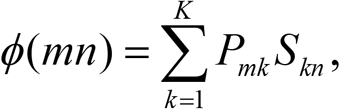

Where the unit *P_mk_S_kn_* represents the influence of mutations of category *m* attributed to signature *k* in tumor *n*. Then the probability effects in mutations of category *m* due to signature *k* in tumor *n* can be expressed as:

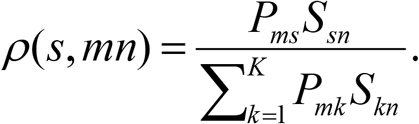

Let us now consider that all genes with the number of mutations of each mutation type in tumor *n* can be calculated in matrix **Γ** that designed as following:

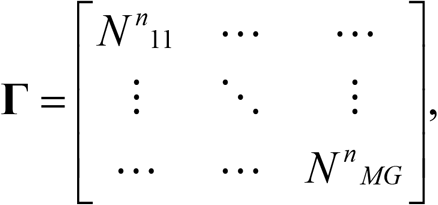

where the *G* represents the number of gene lists in the dataset. Here, we define *ρ^n^_mg_* as the impact factor of mutations of category *m* in sample *n* due to a gene *g*, and it can be calculated by the following formula:

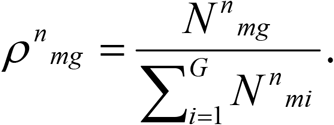

Then the contribution of mutational signature *s* to mutations of category *m* in a gene *g* in tumor *n* can be estimated as:

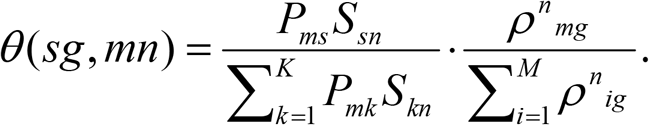

Therefore, we consider that the cumulative abundance of a gene *g* that is attributed to mutational signature *s* in tumor *n* can be estimated as follows:

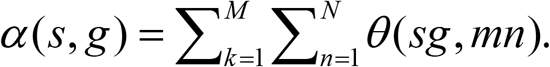

On the other hand, considering the influence of gene length, we also define the relative abundance of a gene *g* that is attributed to mutational signature *s* in tumor *n*, which can be calculated as follows:

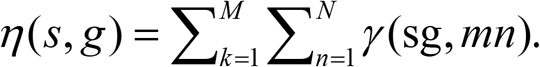

Where *l_g_* represents the length of the gene *g*. And γ(*sg*, *mn*) is on behalf of the relative abundance of mutational signature *s* to mutations of category *m* in a gene *g* in tumor *n* can be estimated as:

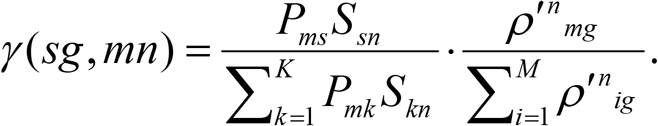

Where the *ρ′^n^_mg_* can be calculated by the following formula:

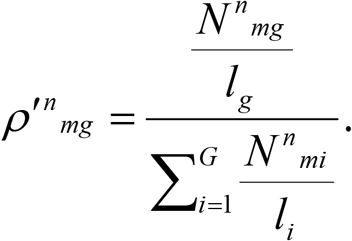

Finally, based on the research results of the previous workers [7, 14, 21, 35], we integrated and optimized some methods to evaluate the association between gene mutation and mutational signatures. We hope that through these methods, we can provide a convenient program interface for researchers, and further provide new ideas for the study of tumor mechanism.

### Prognostic analysis

Kaplan–Meier survival analysis and Cox proportional hazards model were used to analyze an association between cancer-related genes and prognosis. Kaplan–Meier survival and Cox regression analyses were carried out with the R survival package (2.40-1). P-value less than 0.05 was considered to be statistically significant.

## Acknowledgements

This work was funded by a foundation for young creative talents of department education of Guangdong (Natural Science), Grant No. 2019KQNCX067 and science and technology planning project of guangzhou, Grant No. 006259497026.

## Supplementary Figure Text

**Supplementary Figure 1. Comparison of signature contributions identified with deconstructSigs, MutationalPatterns and RNMF framework.** (a) Scatterplots represent the relationship between the weighted proportions calculated using three methods on a set of exon region of 1073 ESCC samples. Each point plotted represents the weights assigned by both methods to one signature detected in a individual. (b) The runtime (elapsed time) in seconds to find the optimal linear combination of mutational signatures from 1073 samples for both packages.

**Supplementary Figure 2. Mutation rate of 508 ESCC.** Lego plot and bar plot representation of mutation patterns in 508 ESCC with whole genome sequencing. Single-nucleotide substitutions are divided into six categories with 16 surrounding flanking bases. Inset pie chart shows the proportion of six categories of mutation patterns. And 83 mutation types were used to display the small insert and deletion. Each column represents the mutation proportion of this mutation type.

**Supplementary Figure 3. Deciphering mutational signatures from a set of mutational catalogs from 508 ESCC.** (a) Identifying the number of SBS signatures in a set of 508 ESCC genomes based on strong stability, low error rate and more stable gradient of error. We choose the classification with stability not less than 0.9 as the best number of SBS signatures for final decomposition, in which the green clipper on each graph indicates the current classification position to be selected. The light purple band represents the confidence region of stability change under the current classification number. (b) The classification selection is adopted by SigProfilerExtractor method, and the gray strip represents the current optimal number of selected signatures. (c) Identifying the number of ID signatures in a set of 508 ESCC genomes based on strong stability, low error rate and more stable gradient of error. We choose the classification with stability not less than 0.9 as the best number of ID signatures for final decomposition, in which the green clipper on each graph indicates the current classification position to be selected. The light purple band represents the confidence region of stability change under the current classification number.

**Supplementary Figure 4. SBS mutational signatures extracted from 508 WGS cases of Chinese ESCC**. The classifications of each mutation type (SBS, 96 classes) are shown in the picture, separately. Each color is used to illustrate the positions of each mutation subtype on each plot.

**Supplementary Figure 5. Heatmap of the cosine similarity between mutational signatures and COSMIC signatures.** (a) The cosine similarity between SBS signatures and COSMIC signatures with version 2 is evaluated, and the extraction methods of mutational signatures include *RNMF* and SigProfilerExtractor. (b) The cosine similarity between SBS signatures which deciphered by SigProfilerExtractor and COSMIC signatures with version 3. (c) Cosine similarity comparison of SBS signatures between *RNMF* and SigProfilerExtractor. The red mark represents the signature with the highest similarity.

**Supplementary Figure 6**. (a) Lego plot representation of mutation patterns in exon region of 1073 ESCC cases. Single-nucleotide substitutions are divided into six categories with 16 surrounding flanking bases. Inset pie chart shows the proportion of six categories of mutation patterns. (b) Based on the background mutation contribution probability of COSMIC Mutational Signatures (v2.0 - March 2015), each color represents a mutational signature, the length of each column represents the contribution proportion of mutation to the signature, the red mark represents the signature most similar to the 12 mutational signature, and the green arrow and green font indicate that this signature is very similar to the 12 mutational signature. (c) The proportion of 1073 ESCC samples in 12 mutational signatures, each color represents a feature, in which red represents the proportion of ubiquitous signatures.

**Supplementary Figure 7**. (a) Association of CCA of cancer-related genes from the COSMIC census assigned to SBS signatures with prognosis. Kaplan-Meier survival analysis classified by the status that CCA of genes assigned to SBS signatures with a threshold 6%. (b) Multivariate Cox regression analysis of CCA of genes assigned to SBS signatures with age, gender, stage and CCA of genes assigned to SBS signatures. Red is marked with significant items.

**Supplementary Figure 8**. (a) Box plot showing that the SBS signatures were associated with cancer-related genes mutations (only SNV), where n represents the number of samples. (b-d) CCA enrichment analysis and mutational signature enrichment analyses identify an association between somatic mutations and activity of signatures in ESCC. Here, we use two datasets: exon regions of 1073 and 508 ESCC cases. First, the median CCA of each gene in the current mutational signature is calculated, and then the contribution importance of each gene is calculated by PERMUTATION test or Fisher’s test to study the association between gene and feature. The regular tumors in the figure represent the samples with non hypermutated. For genes mutated in >5% of samples, the CCA of genes attributed to SBS or ID signatures was compared in tumors with wild-type versus mutated copies of the gene. Genes with FDR q < 0.1 are highlighted in red. (e) Association of *TP53* mutations with prognosis. Kaplan-Meier survival analysis classified by the status that *TP53* hotspot mutation (p.R248Q). (f) Multivariate Cox regression analysis of *TP53* CCA assigned to SBS1 with age, gender, stage, Location, *TP53* hotspot mutation p.R248Q status and CCA of *TP53* assigned to SBS1*. (g) CCA enrichment analysis and mutational signature enrichment analyses identify an association between somatic mutations and activity of signatures in ESCC. First, the median CCA of each gene in the current mutational signature is calculated, and then the contribution importance of each gene is calculated by PERMUTATION test or Fisher’s test to study the association between gene and feature. The regular tumors in the figure represent the samples with non hypermutated. For genes mutated in >5% of samples, the CCA of genes attributed to SBS or ID signatures was compared in tumors with wild-type versus mutated copies of the gene. Genes with FDR q < 0.1 are highlighted in red.

**Supplementary Figure 9. The application and association of CCA matrix of genes.** (a) Accuracy evaluation of model test between G2 group and G4 group. Wilcoxon rank sum test with two-sided was used for statistical significance. (b) Full schematic representation of DenseNet-121. (c) The model DenseNet-121 was trained 10 times for each sub-feature data of G2-G4 and G2G3-G4. For each result, we randomly select 90% of the samples as training, and the remaining 10% as test data set for analysis. Each box represents a mutational signature and is marked in a different color. One way ANOVA test was used for statistical significance. The higher accuracy level of mutational signature is marked in red. (d) The distribution of G3 groups in the parameter model obtained by G2-G4. A color represents a group, a curve represents a density curve. (e) The difference of sample contribution in different groups was compared. Wilcoxon rank sum test with two-sided was used for statistical significance.

## References

[1] Stratton M R, Campbell P J, Futreal P A. The cancer genome. Nature 458, 719–724 (2009).

[2] Pfeifer G P. Environmental exposures and mutational patterns of cancer genomes. Genome medicine 2, 54–57 (2010).

[3] Boland C R, Goel A. Microsatellite instability in colorectal cancer. Gastroenterology 138, 2073–2087 (2010).

[4] Nik-Zainal S, Alexandrov L B, Wedge D C, et al. Mutational processes molding the genomes of 21 breast cancers. Cell 149, 979–993 (2012).

[5] Alexandrov L B, Stratton M R. Mutational signatures: the patterns of somatic mutations hidden in cancer genomes. Current opinion in genetics & development 24, 52–60 (2014).

[6] Stratton M R. Exploring the genomes of cancer cells: progress and promise. Science 331, 1553–1558 (2011).

[7] Letouzé E, Shinde J, Renault V, et al. Mutational signatures reveal the dynamic interplay of risk factors and cellular processes during liver tumorigenesis. Nature communications 8, 1–13 (2017).

[8] Davies H, Glodzik D, Morganella S, et al. HRDetect is a predictor of BRCA1 and BRCA2 deficiency based on mutational signatures. Nature Medicine 23, 517–525 (2017).

[9] Ma J, Setton J, Lee N Y, et al. The therapeutic significance of mutational signatures from DNA repair deficiency in cancer. Nature communications 9, 1–12 (2018).

[10] Wang S, Jia M, He Z, et al. APOBEC3B and APOBEC mutational signature as potential predictive markers for immunotherapy response in non-small cell lung cancer. Oncogene 37, 3924–3936 (2018).

[11] Harris R S. Cancer mutation signatures, DNA damage mechanisms, and potential clinical implications. Genome medicine, 2013, 5(9): 87.

[12] Wagener R, Alexandrov L B, Montesinos-Rongen M, et al. Analysis of mutational signatures in exomes from B-cell lymphoma cell lines suggest APOBEC3 family members to be involved in the pathogenesis of primary effusion lymphoma. Leukemia 29, 1612–1615 (2015).

[13] Helleday T, Eshtad S, Nik-Zainal S. Mechanisms underlying mutational signatures in human cancers. Nature Reviews Genetics 15, 585–598 (2014).

[14] Kim J, Mouw K W, Polak P, et al. Somatic ERCC2 mutations are associated with a distinct genomic signature in urothelial tumors. Nature genetics 48, 600–606 (2016).

[15] Alexandrov L B, Nik-Zainal S, Wedge D C, et al. Deciphering signatures of mutational processes operative in human cancer. Cell reports 3, 246–259 (2013).

[16] Gehring J S, Fischer B, Lawrence M, et al. SomaticSignatures: inferring mutational signatures from single-nucleotide variants. Bioinformatics 31, 3673–3675 (2015).

[17] Shiraishi Y, Tremmel G, Miyano S, et al. A simple model-based approach to inferring and visualizing cancer mutation signatures. PLoS genetics 11, 1005657_1–21 (2015).

[18] Rosales R A, Drummond R D, Valieris R, et al. signeR: an empirical Bayesian approach to mutational signature discovery. Bioinformatics 33, 8–16 (2017).

[19] Ardin M, Cahais V, Castells X, et al. MutSpec: a Galaxy toolbox for streamlined analyses of somatic mutation spectra in human and mouse cancer genomes. BMC bioinformatics 17, 170_1–10 (2016).

[20] Blokzijl F, Janssen R, van Boxtel R, et al. MutationalPatterns: comprehensive genome-wide analysis of mutational processes. Genome medicine 10, 33_1–11 (2018).

[21] Xing R, Zhou Y, Yu J, et al. Whole-genome sequencing reveals novel tandem-duplication hotspots and a prognostic mutational signature in gastric cancer. Nature communications 10, 1–13 (2019).

[22] Baez-Ortega A, Gori K. Computational approaches for discovery of mutational signatures in cancer. Briefings in bioinformatics 20, 77–88 (2019).

[23] Alexandrov L B, Kim J, Haradhvala N J, et al. The repertoire of mutational signatures in human cancer. Nature 578, 94–101(2020).

[24] Alexandrov L B, Nik-Zainal S, Wedge D C, et al. Signatures of mutational processes in human cancer. Nature 500, 415–421 (2013).

[25] Poon S L, Pang S T, McPherson J R, et al. Genome-wide mutational signatures of aristolochic acid and its application as a screening tool. Science translational medicine 5, 197ra101 (2013).

[26] Poon S L, Huang M N, Choo Y, et al. Mutation signatures implicate aristolochic acid in bladder cancer development. Genome medicine 7, 1–10 (2015).

[27] Schulze K, Imbeaud S, Letouzé E, et al. Exome sequencing of hepatocellular carcinomas identifies new mutational signatures and potential therapeutic targets. Nature genetics 47, 505–511 (2015).

[28] Merlevede J, Droin N, Qin T, et al. Mutation allele burden remains unchanged in chronic myelomonocytic leukaemia responding to hypomethylating agents. Nature communications 7, 1–13 (2016).

[29] Nik-Zainal S, Davies H, Staaf J, et al. Landscape of somatic mutations in 560 breast cancer whole-genome sequences. Nature 534, 47–54 (2016).

[30] Petljak M, Alexandrov L B. Understanding mutagenesis through delineation of mutational signatures in human cancer. Carcinogenesis 37, 531–540 (2016).

[31] Polak P, Kim J, Braunstein L Z, et al. A mutational signature reveals alterations underlying deficient homologous recombination repair in breast cancer. Nature genetics 49, 1476–1486 (2017).

[32] Fischer A, Illingworth C J R, Campbell P J, et al. EMu: probabilistic inference of mutational processes and their localization in the cancer genome. Genome biology 14, 1–10 (2013).

[33] Rosenthal R, McGranahan N, Herrero J, et al. DeconstructSigs: delineating mutational processes in single tumors distinguishes DNA repair deficiencies and patterns of carcinoma evolution. Genome biology 17, 1–11 (2016).

[34] Ramos A H, Lichtenstein L, Gupta M, et al. Oncotator: cancer variant annotation tool. Human mutation 36, 2423–2429 (2015).

[35] Lin D C, Dinh H Q, Xie J J, et al. Identification of distinct mutational patterns and new driver genes in oesophageal squamous cell carcinomas and adenocarcinomas. Gut 67, 1769–1779 (2018).

[36] Chen W, Zheng R, Baade PD et al. Cancer Statistics in China, CA: A Cancer Journal for Clinicians 66, 115–132 (2016).

[37] Cui Y, Chen H, Xi R, et al. Whole-genome sequencing of 508 patients identifies key molecular features associated with poor prognosis in esophageal squamous cell carcinoma. Cell research 30, 902–913 (2020).

[38] Burns, M.B., Temiz, N.A., Harris, R.S.. Evidence for APOBEC3B mutagenesis in multiple human cancers. Nat. Genet. 45, 977–983 (2013).

[39] Roberts, S.A., Lawrence, et al. An APOBEC cytidine deaminase mutagenesis pattern is widespread in human cancers. Nat. Genet. 45, 970–976 (2013).

[40] Zhang L, Zhou Y, Cheng C, et al. Genomic analyses reveal mutational signatures and frequently altered genes in esophageal squamous cell carcinoma. The American Journal of Human Genetics 96, 597–611 (2015).

[41] Degasperi, A., Amarante, T.D., Czarnecki, J. et al. A practical framework and online tool for mutational signature analyses show intertissue variation and driver dependencies. Nat Cancer 1, 249–263 (2020).

[42] Giacomelli, A.O., Yang, X., Lintner, R.E. et al. Mutational processes shape the landscape of TP53 mutations in human cancer. Nat Genet 50, 1381–1387 (2018).

[43] Degasperi, A., Amarante, T.D., Czarnecki, J. et al. A practical framework and online tool for mutational signature analyses show intertissue variation and driver dependencies. Nat Cancer 1, 249–263 (2020).

[44] X.C. Li, M.Y. Wang, M. Yang, et al. A mutational signature associated with alcohol consumption and prognostically significantly mutated driver genes in esophageal squamous cell carcinoma. Annals of Oncology 29, 938–944 (2018).

[45] Ketkar N. Introduction to pytorch. Deep learning with python. Apress, Berkeley, CA, 195–208 (2017).

[46] He K, Zhang X, Ren S, et al. Deep residual learning for image recognition. Proceedings of the IEEE conference on computer vision and pattern recognition. 770–778 (2016).

[47] Huang G, Liu Z, Van Der Maaten L, et al. Densely connected convolutional networks. Proceedings of the IEEE conference on computer vision and pattern recognition. 4700–4708 (2017).

[48] Pleiss G, Chen D, Huang G, et al. Memory-efficient implementation of densenets. arXiv preprint arXiv:1707.06990 (2017).

[49] Szegedy C, Ioffe S, Vanhoucke V, et al. Inception-v4, inception-resnet and the impact of residual connections on learning. Proceedings of the AAAI Conference on Artificial Intelligence 31, (2017).

[50] Sandler M, Howard A, Zhu M, et al. Mobilenetv2: Inverted residuals and linear bottlenecks. Proceedings of the IEEE conference on computer vision and pattern recognition, 4510–4520 (2018).

[51] Howard A, Sandler M, Chu G, et al. Searching for mobilenetv3. Proceedings of the IEEE/CVF International Conference on Computer Vision. 1314–1324 (2019).

[52] Bottou L. Large-scale machine learning with stochastic gradient descent. Proceedings of COMPSTAT’2010. Physica-Verlag HD, 177–186 (2010).

[53] Steven A. Roberts, Dmitry A. Gordenin. Clustered and genome-wide transient mutagenesis in human cancers: Hypermutation without permanent mutators or loss of fitness. BioEssays 36,382–393 (2014).

[54] Alexandrov, L., Jones, P., Wedge, D. et al. Clock-like mutational processes in human somatic cells. Nat Genet 47, 1402–1407 (2015).

[55] Zhang J, Wei L, Feng X, et al. Pattern expression nonnegative matrix factorization: algorithm and applications to blind source separation. Computational intelligence and neuroscience 2008, 1–10 (2008).

[56] Rajabi R, Ghassemian H. Spectral unmixing of hyperspectral imagery using multilayer NMF. IEEE Geoscience and Remote Sensing Letters 12, 38–42 (2014).

[57] Alsabti K, Ranka S, Singh V. An efficient k-means clustering algorithm. Syracuse University, (1997).

