## Supplemental files Figure for "A practical framework RNMF for the potential mechanism of cancer progression with the analysis of genes cumulative contribution abundance": Sup Fig. 1 .pdf

a

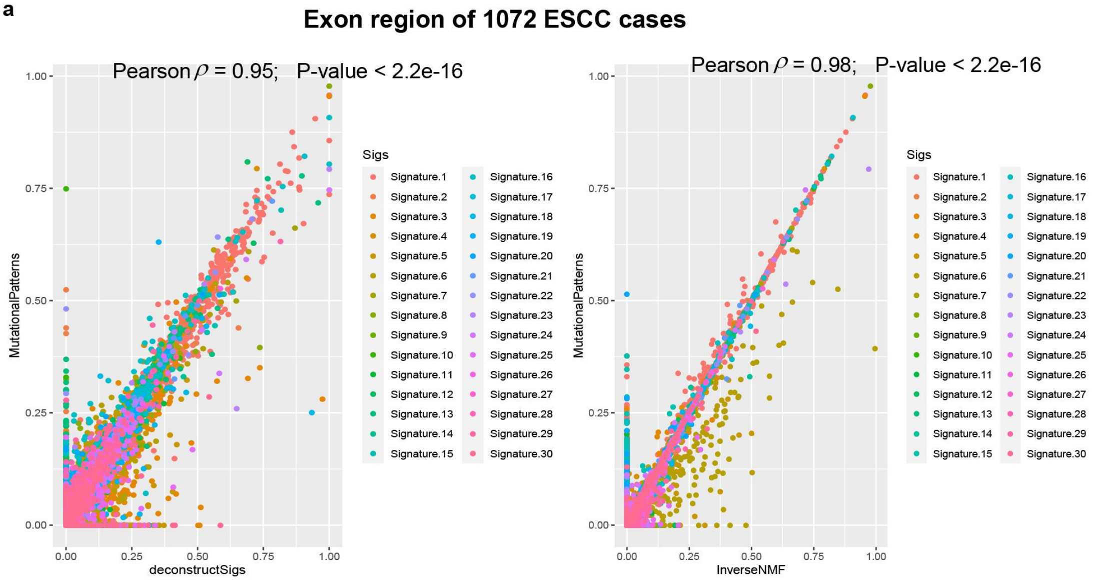


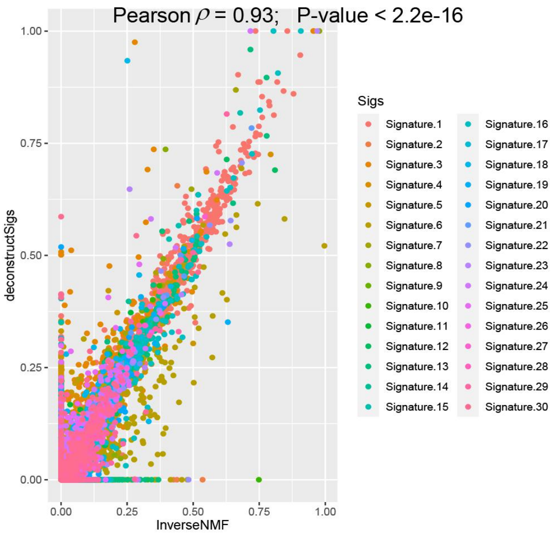


b

| | InverseNMF | MutationalPatterns | deconstructSigs |
|---|---|---|---|
| Runtime (1072 samples) | 44.9 s | 22.8 s | 9 min 5.5 s |

**Supplementary figure 5. Comparison of signature contributions identified with deconstructSigs, MutationalPatterns and RNMF framework.** (a) Scatterplots represent the relationship between the weighted proportions calculated using three methods on a set of exon region of 1072 ESCC samples. Each point plotted represents the weights assigned by both methods to one signature detected in a individual. (b) The runtime (elapsed time) in seconds to find the optimal linear combination of mutational signatures from 1072 samples for both packages.
