## Supplemental files Figure for "A practical framework RNMF for the potential mechanism of cancer progression with the analysis of genes cumulative contribution abundance": Sup Fig. 7 .pdf

a

Survival analysis

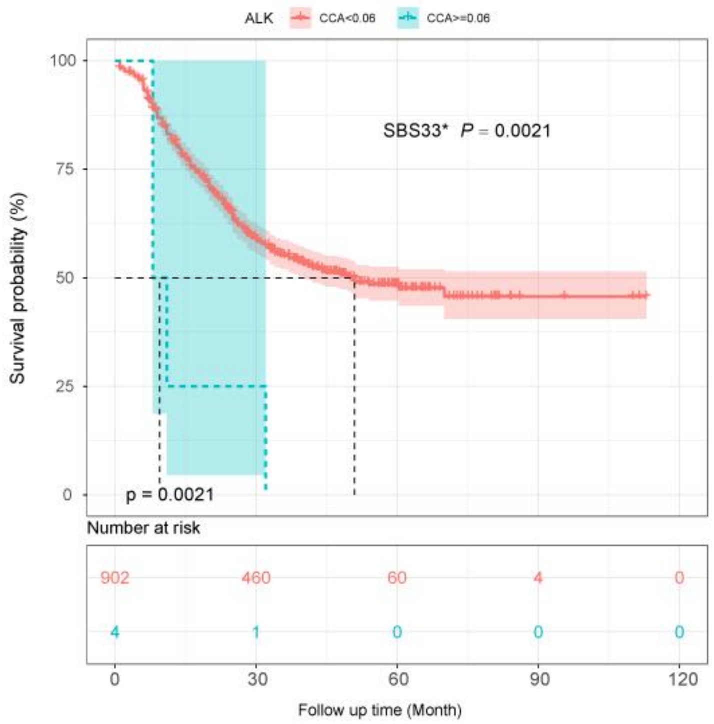


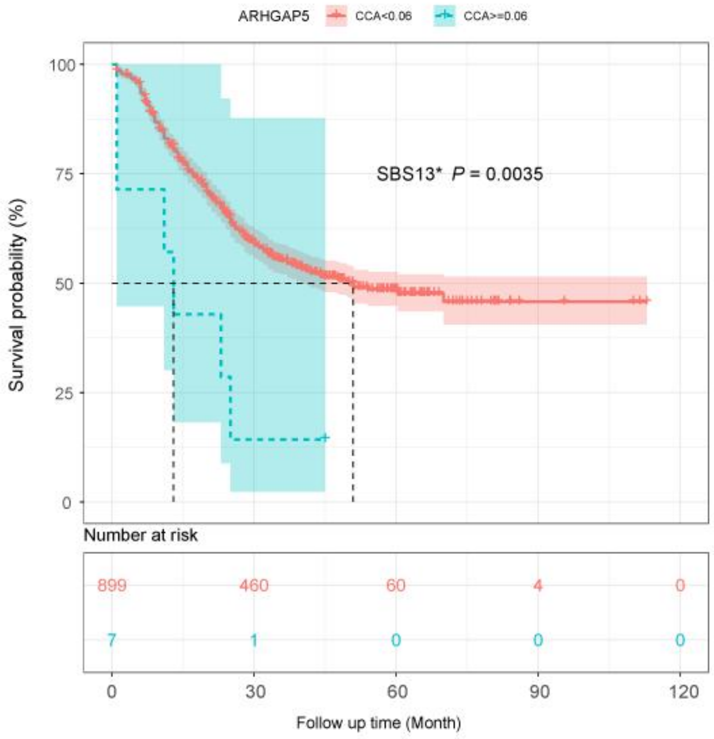

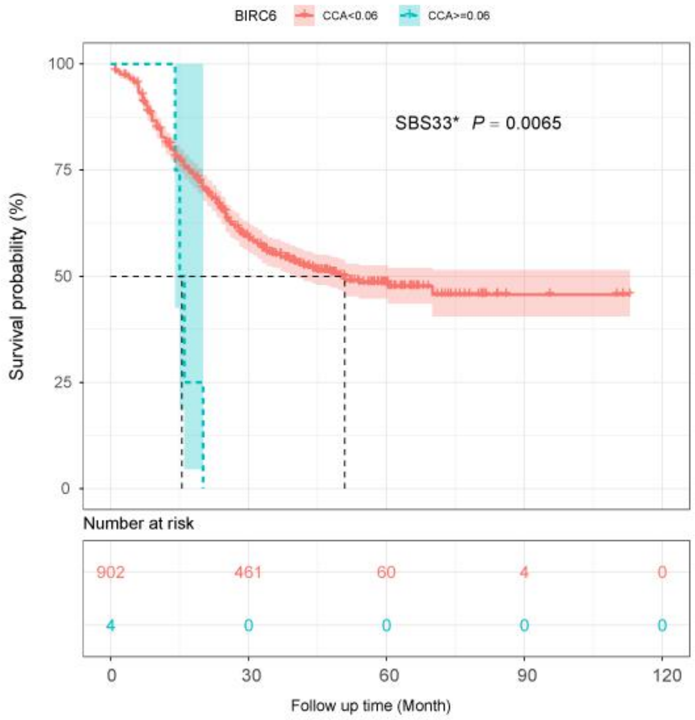
BIRC6
CCA<0.06
CCA>=0.06
SBS33* P = 0.0065
Survival probability (%)
100
75
50
25
0
Number at risk
902
461
60
4
0
4
0
0
0
0
0
30
60
90
120
Follow up time (Month)

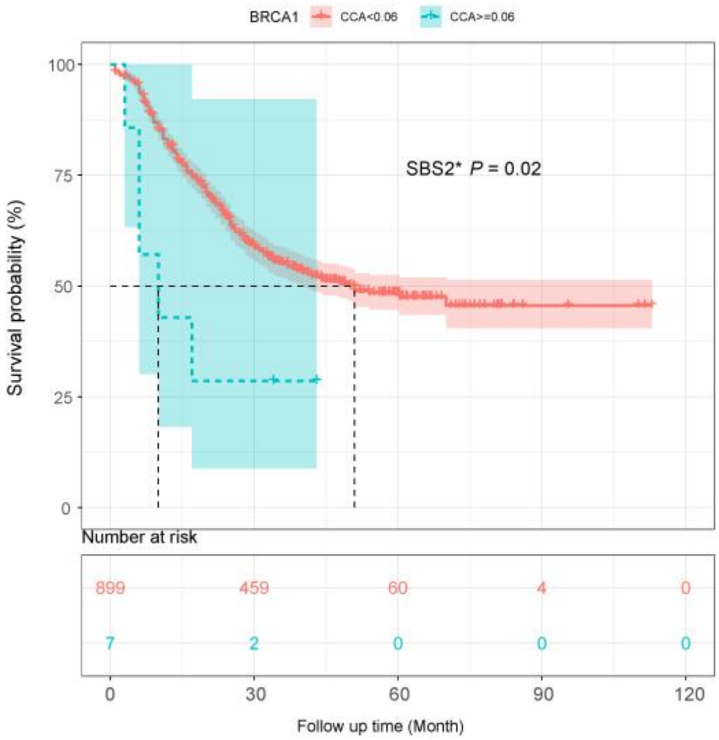
BRCA1
CCA<0.06
CCA>=0.06
SBS2* P = 0.02
Survival probability (%)
100
75
50
25
0
Number at risk
899
459
60
4
0
7
2
0
0
0
0
30
60
90
120
Follow up time (Month)

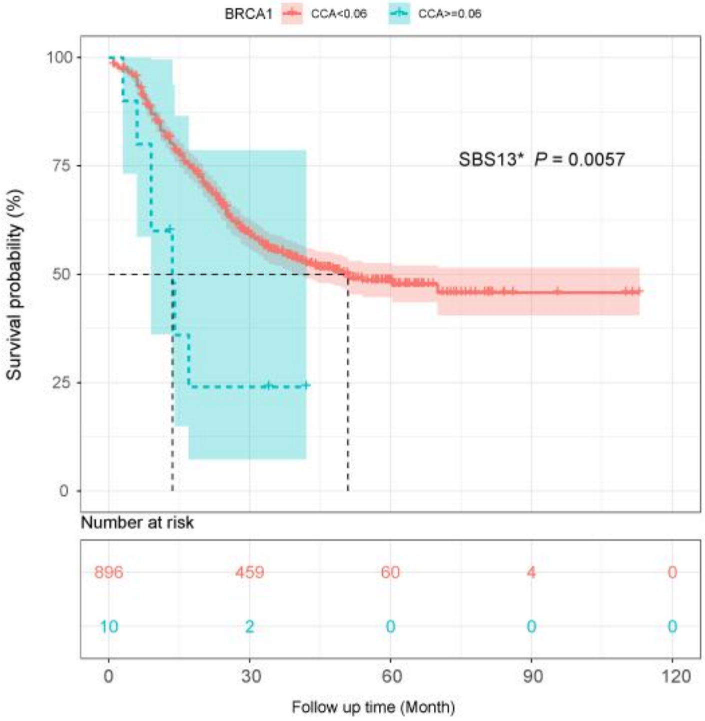
BRCA1
CCA<0.06
CCA>=0.06
SBS13* P = 0.0057
Survival probability (%)
100
75
50
25
0
Number at risk
896
459
60
4
0
10
2
0
0
0
0
30
60
90
120
Follow up time (Month)

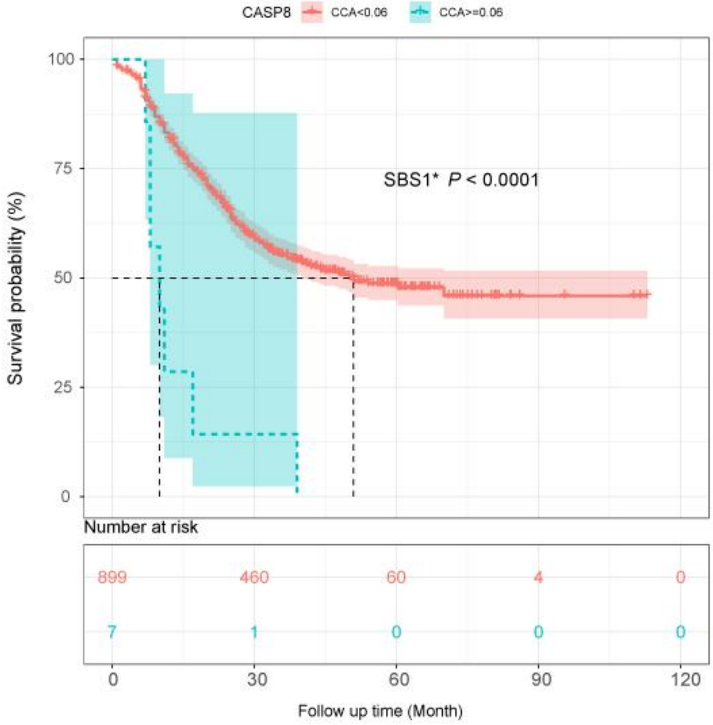
CASP8
CCA<0.06
CCA>=0.06
SBS1* P < 0.0001
Survival probability (%)
100
75
50
25
0
Number at risk
899
460
60
4
0
7
1
0
0
0
0
30
60
90
120
Follow up time (Month)

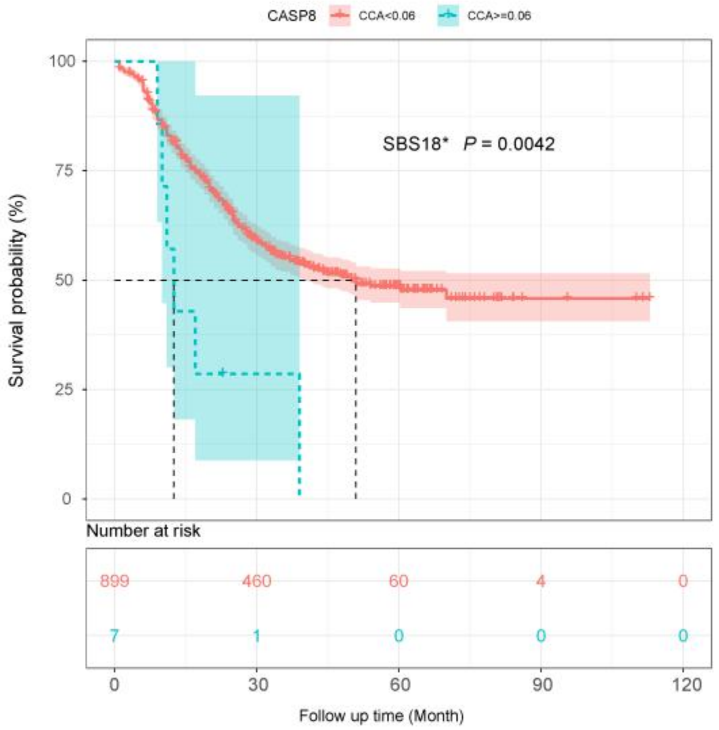
CASP8
CCA<0.06
CCA>=0.06
SBS18* P = 0.0042
Survival probability (%)
100
75
50
25
0
Number at risk
899
460
60
4
0
7
1
0
0
0
0
30
60
90
120
Follow up time (Month)

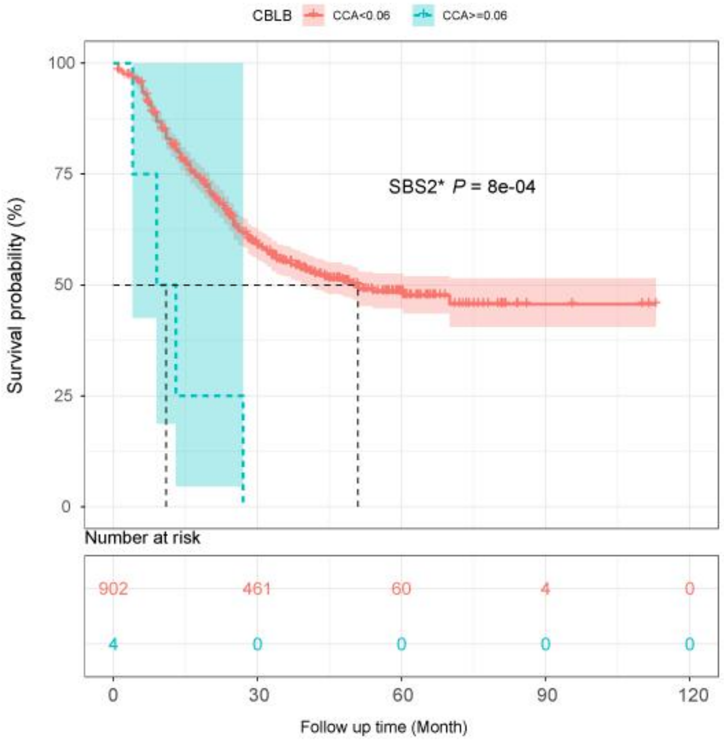
CBLB
CCA<0.06
CCA>=0.06
SBS2* P = 8e-04
Survival probability (%)
100
75
50
25
0
Number at risk
902
461
60
4
0
4
0
0
0
0
0
30
60
90
120
Follow up time (Month)

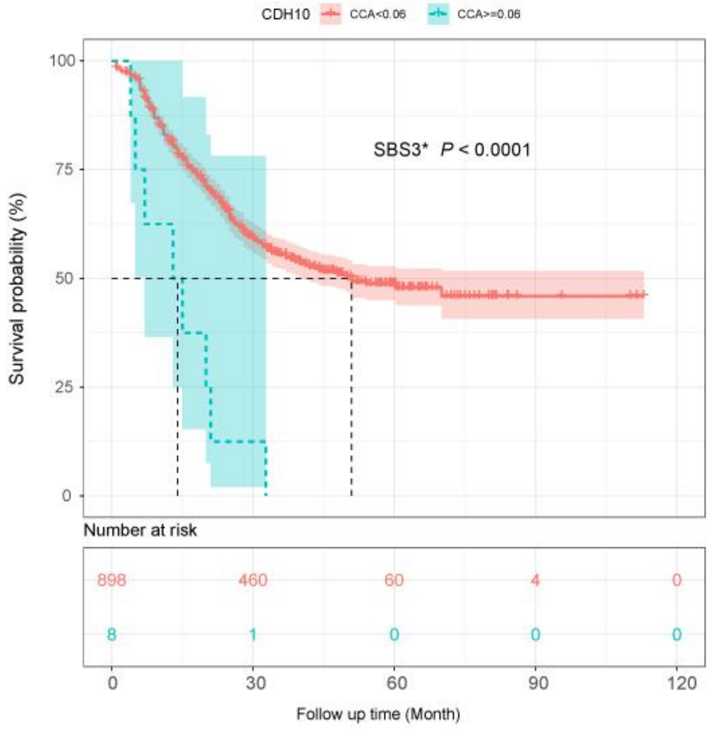
CDH10
CCA<0.06
CCA>=0.06
SBS3* P < 0.0001
Survival probability (%)
100
75
50
25
0
Number at risk
898
460
60
4
0
8
1
0
0
0
0
30
60
90
120
Follow up time (Month)

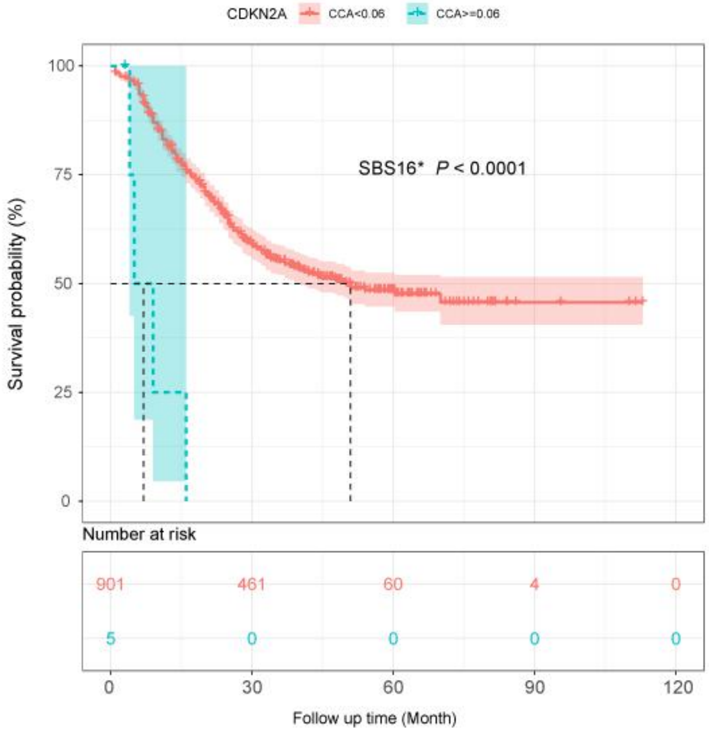
CDKN2A
CCA<0.06
CCA>=0.06
SBS16* P < 0.0001
Survival probability (%)
100
75
50
25
0
Number at risk
901
461
60
4
0
5
0
0
0
0
0
30
60
90
120
Follow up time (Month)

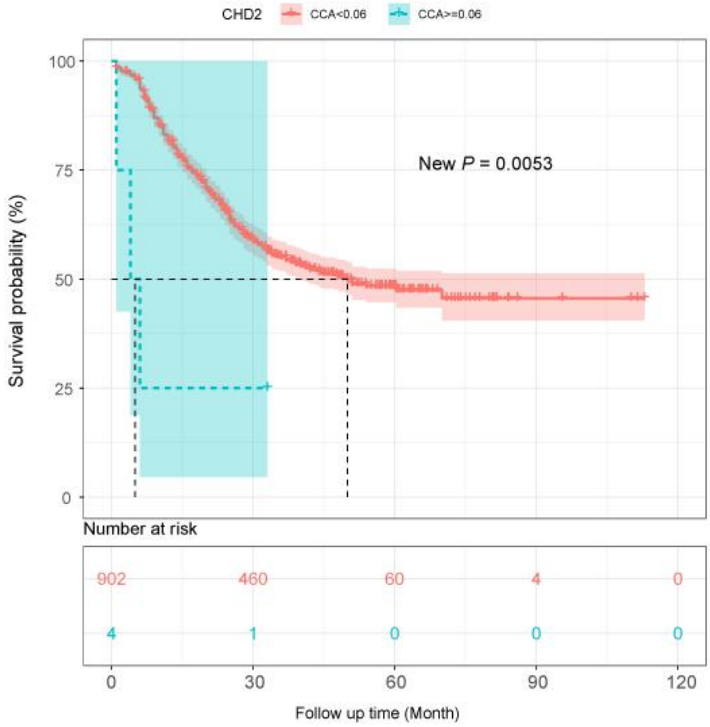
CHD2 CCA<0.06 CCA>=0.06
New P = 0.0053
Survival probability (%)
100
75
50
25
0
Number at risk
902 460 60 4 0
4 1 0 0 0
0 30 60 90 120
Follow up time (Month)

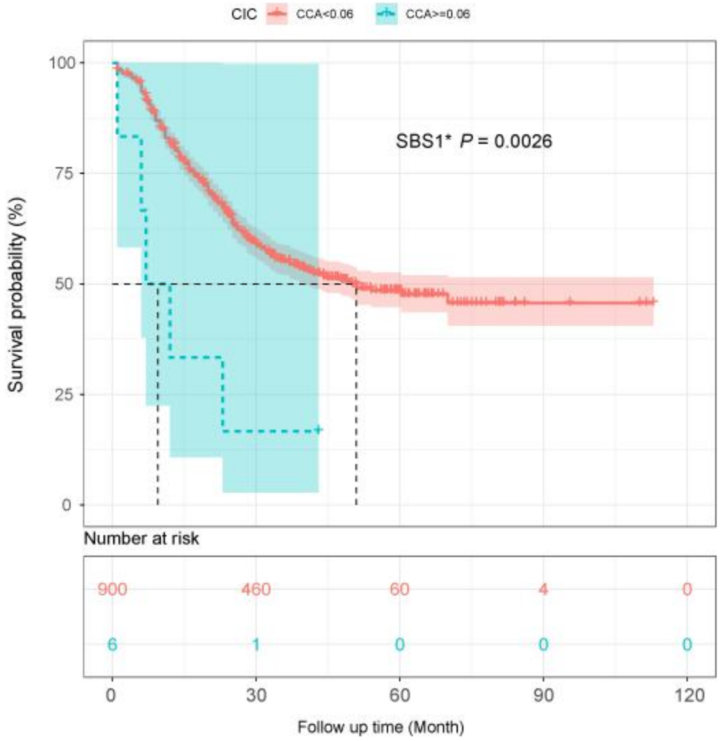
CIC CCA<0.06 CCA>=0.06
SBS1* P = 0.0026
Survival probability (%)
100
75
50
25
0
Number at risk
900 460 60 4 0
6 1 0 0 0
0 30 60 90 120
Follow up time (Month)

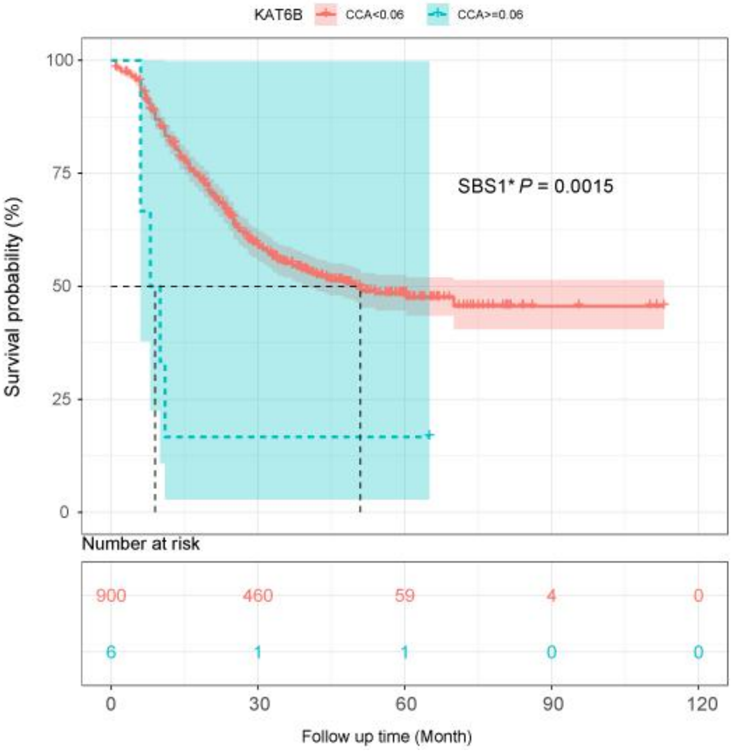
KAT6B
CCA<0.06
CCA>=0.06
SBS1* P = 0.0015
Survival probability (%)
100
75
50
25
0
Number at risk
900
460
59
4
0
6
1
1
0
0
0
30
60
90
120
Follow up time (Month)

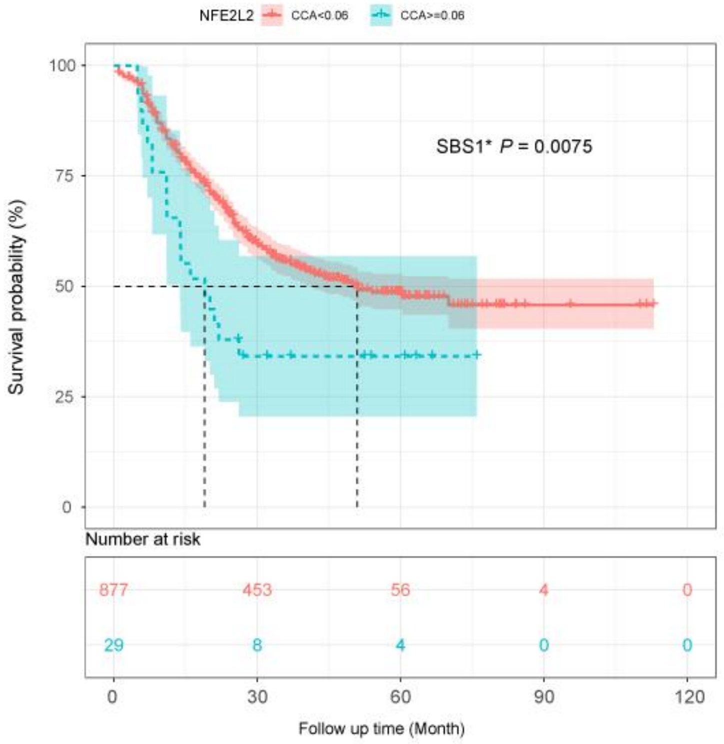
NFE2L2
CCA<0.06
CCA>=0.06
SBS1* P = 0.0075
Survival probability (%)
100
75
50
25
0
Number at risk
877
453
56
4
0
29
8
4
0
0
0
30
60
90
120
Follow up time (Month)

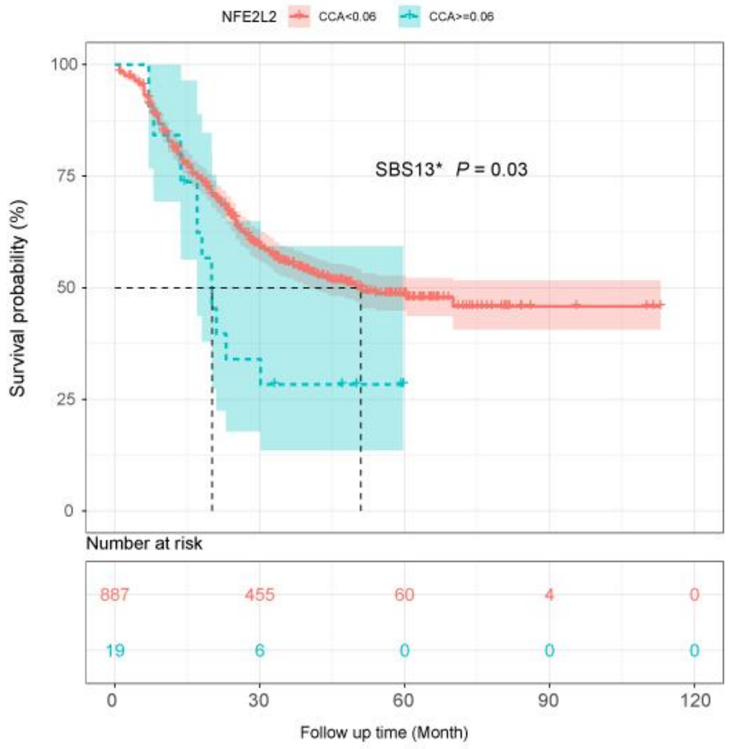
NFE2L2
CCA<0.06
CCA>=0.06
SBS13* P = 0.03
Survival probability (%)
100
75
50
25
0
Number at risk
887
455
60
4
0
19
6
0
0
0
0
30
60
90
120
Follow up time (Month)

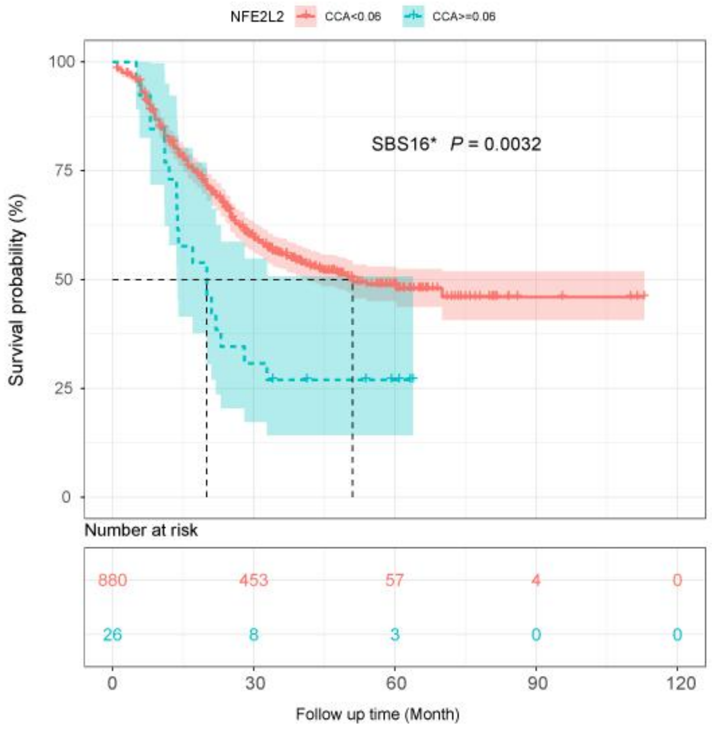
NFE2L2
CCA<0.06
CCA>=0.06
SBS16* P = 0.0032
Survival probability (%)
100
75
50
25
0
Number at risk
880
453
57
4
0
26
8
3
0
0
0
30
60
90
120
Follow up time (Month)

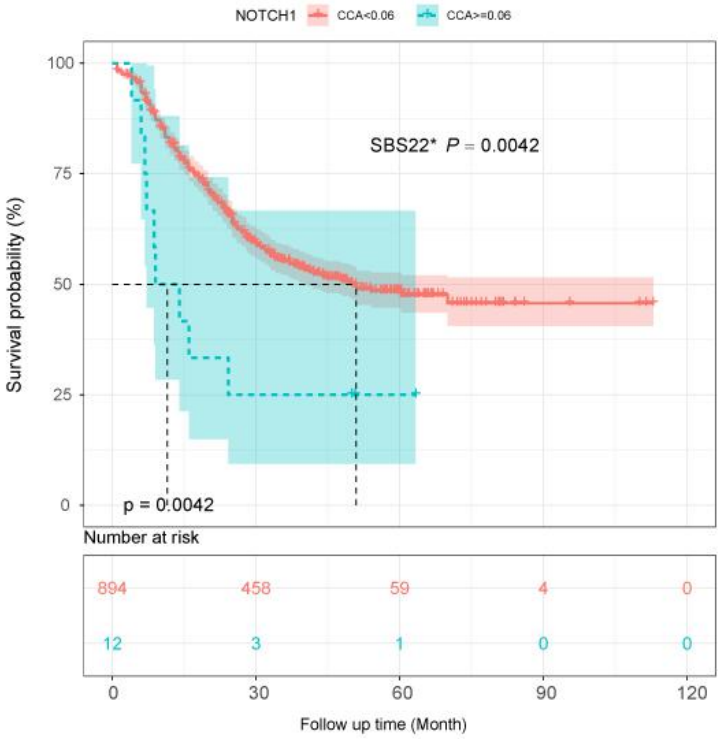

NOTCH1
CCA<0.06
CCA>=0.06
SBS22* P = 0.0042
p = 0.0042
Survival probability (%)
100
75
50
25
0
Number at risk
894
458
59
4
0
12
3
1
0
0
0
30
60
90
120
Follow up time (Month)


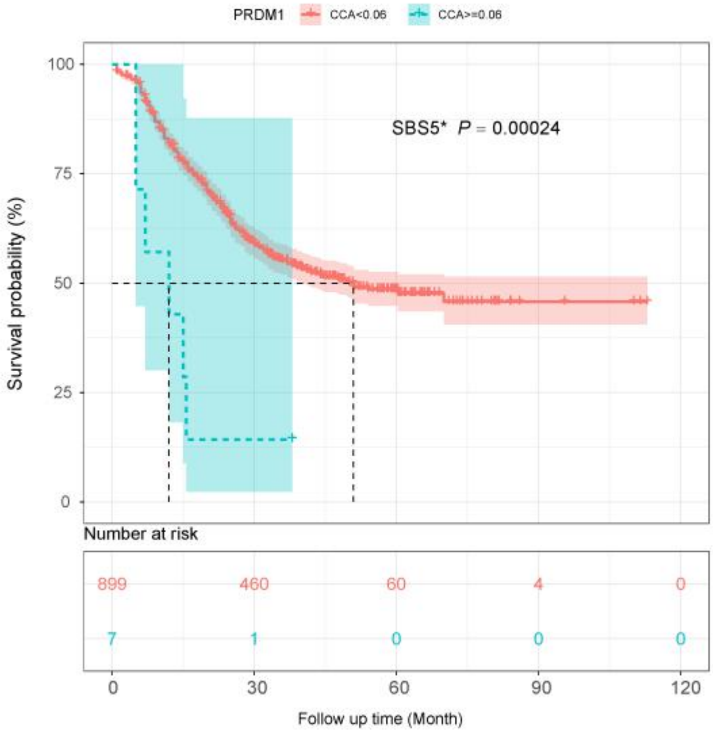

PRDM1
CCA<0.06
CCA>=0.06
SBS5* P = 0.00024
Survival probability (%)
100
75
50
25
0
Number at risk
899
460
60
4
0
7
1
0
0
0
0
30
60
90
120
Follow up time (Month)

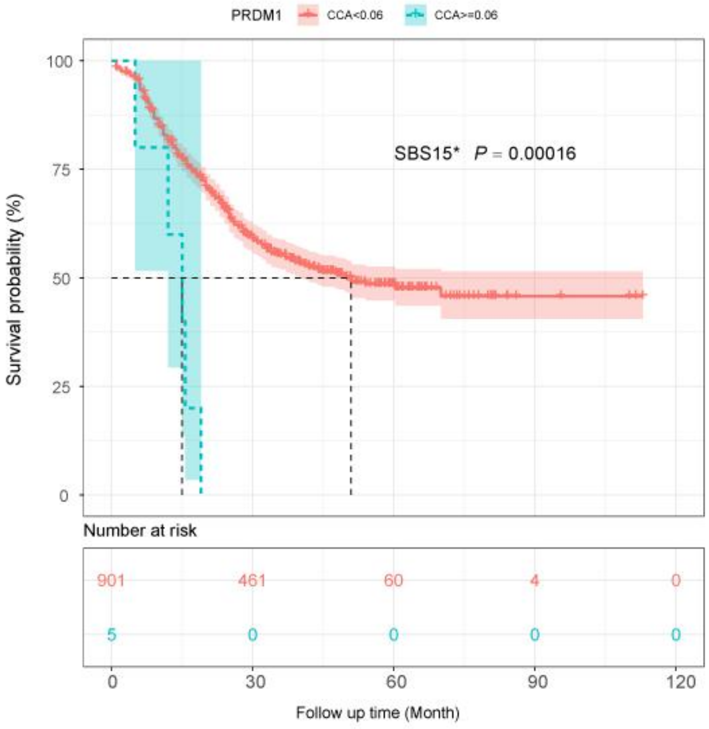
PRDM1
CCA<0.06
CCA>=0.06
SBS15* P = 0.00016
Survival probability (%)
100
75
50
25
0
Number at risk
901 461 60 4 0
5 0 0 0 0
0 30 60 90 120
Follow up time (Month)

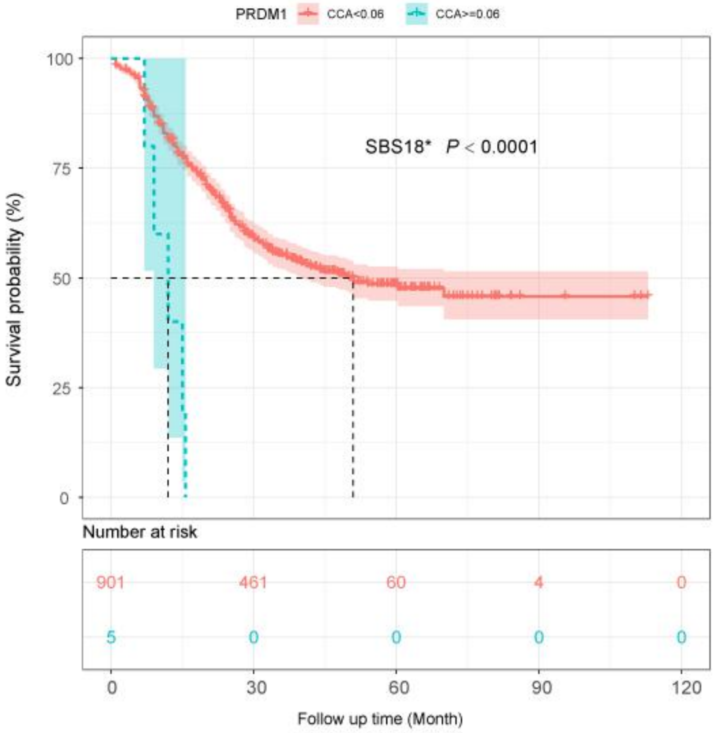
PRDM1
CCA<0.06
CCA>=0.06
SBS18* P < 0.0001
Survival probability (%)
100
75
50
25
0
Number at risk
901 461 60 4 0
5 0 0 0 0
0 30 60 90 120
Follow up time (Month)

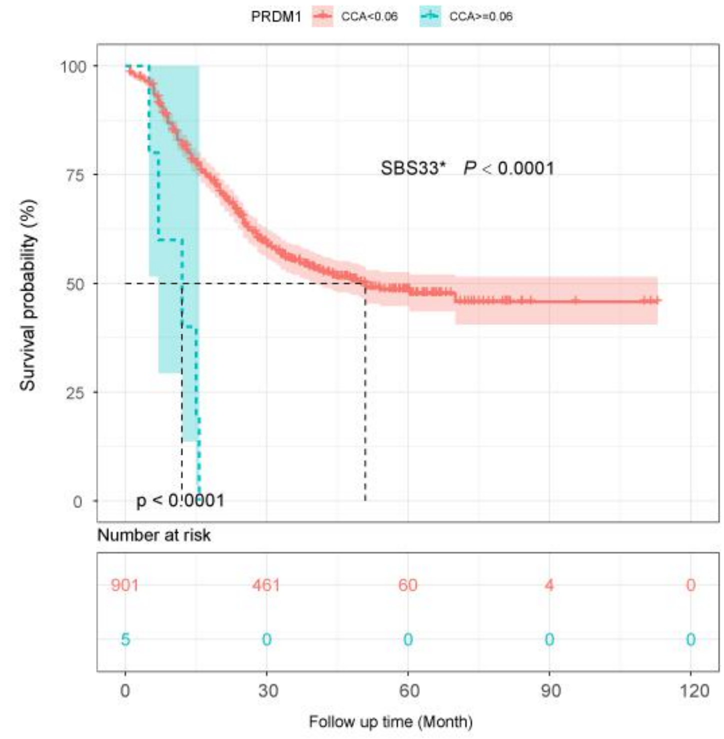
PRDM1
CCA<0.06
CCA>=0.06
SBS33* P < 0.0001
Survival probability (%)
100
75
50
25
0
p < 0.0001
Number at risk
901
461
60
4
0
5
0
0
0
0
0
30
60
90
120
Follow up time (Month)

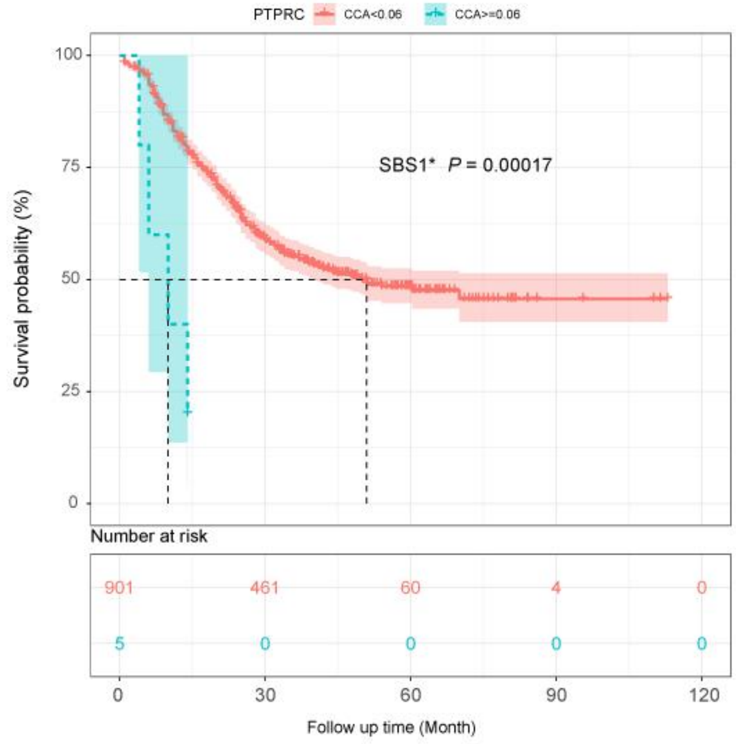
PTPRC
CCA<0.06
CCA>=0.06
SBS1* P = 0.00017
Survival probability (%)
100
75
50
25
0
Number at risk
901
461
60
4
0
5
0
0
0
0
0
30
60
90
120
Follow up time (Month)

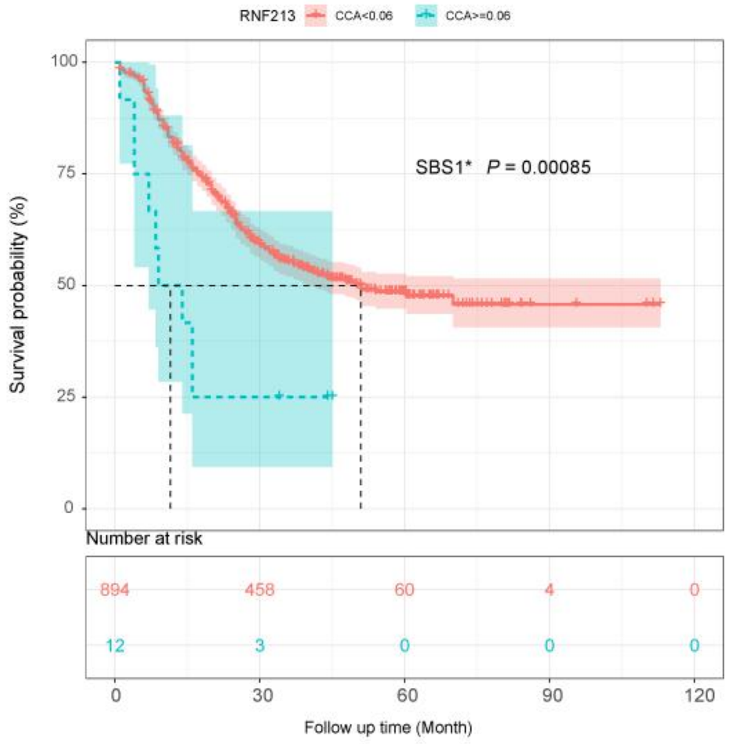
RNF213
CCA<0.06
CCA>=0.06
SBS1* P = 0.00085
Survival probability (%)
100
75
50
25
0
Number at risk
894
458
60
4
0
12
3
0
0
0
0
30
60
90
120
Follow up time (Month)

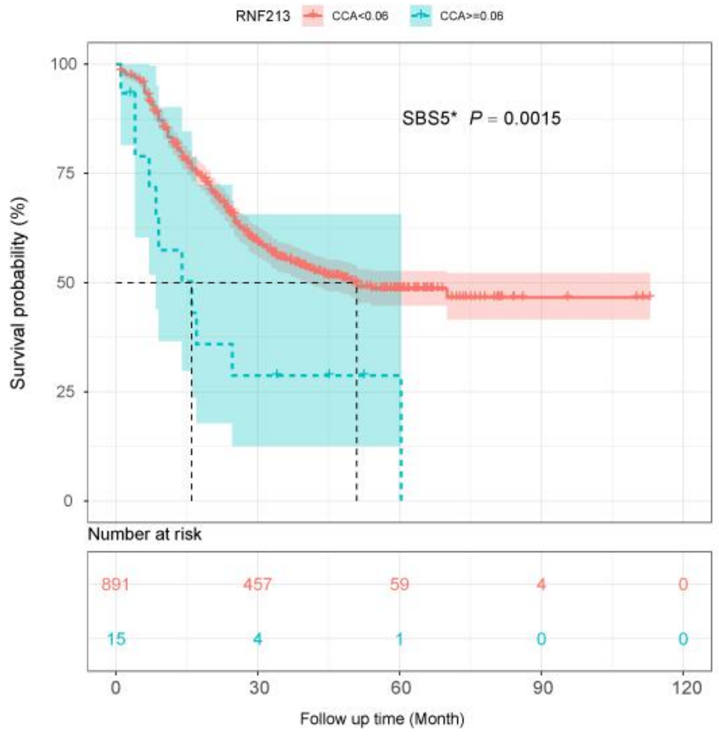
RNF213
CCA<0.06
CCA>=0.06
SBS5* P = 0.0015
Survival probability (%)
100
75
50
25
0
Number at risk
891
457
59
4
0
15
4
1
0
0
0
30
60
90
120
Follow up time (Month)

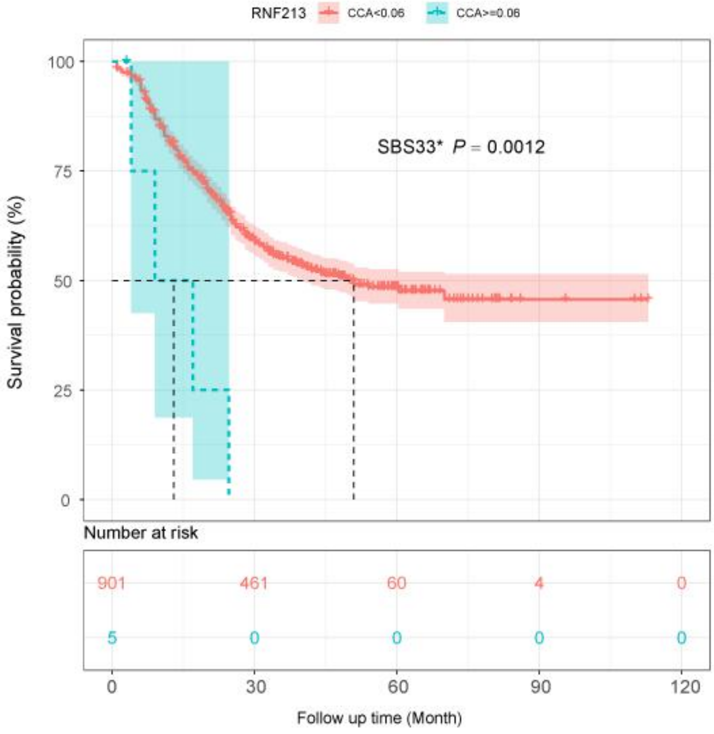
RNF213
CCA<0.06
CCA>=0.06
SBS33* P = 0.0012
Survival probability (%)
100
75
50
25
0
Number at risk
901
461
60
4
0
5
0
0
0
0
0
30
60
90
120
Follow up time (Month)

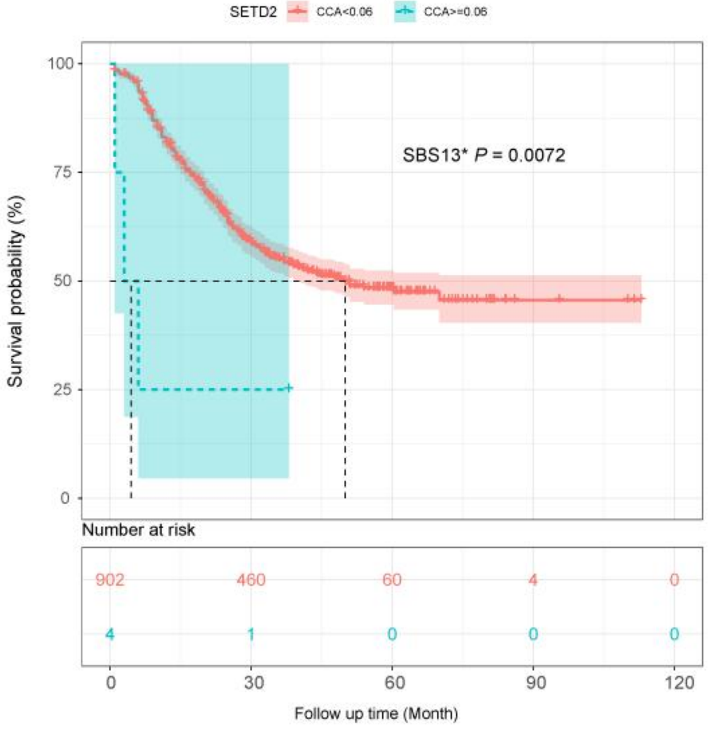
SETD2
CCA<0.06
CCA>=0.06
SBS13* P = 0.0072
Survival probability (%)
100
75
50
25
0
Number at risk
902
460
60
4
0
4
1
0
0
0
0
30
60
90
120
Follow up time (Month)

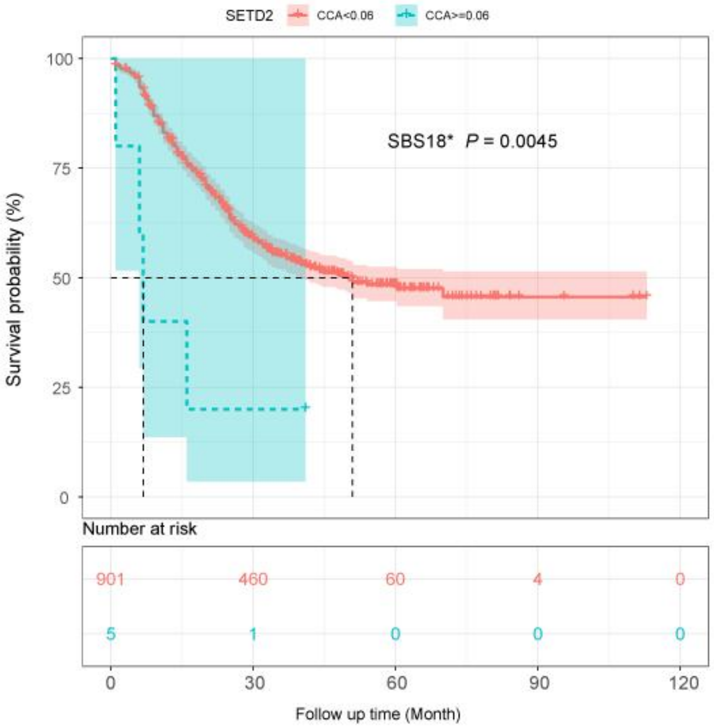
SETD2
CCA<0.06
CCA>=0.06
SBS18* P = 0.0045
Survival probability (%)
100
75
50
25
0
Number at risk
901
460
60
4
0
5
1
0
0
0
0
30
60
90
120
Follow up time (Month)

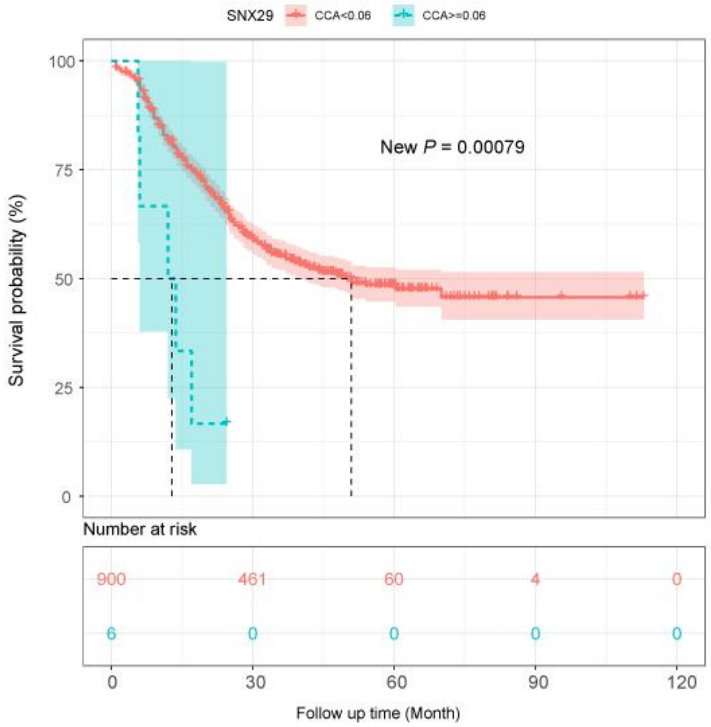
SNX29
CCA<0.06
CCA>=0.06
New P = 0.00079
Survival probability (%)
100
75
50
25
0
Number at risk
900
461
60
4
0
6
0
0
0
0
0
30
60
90
120
Follow up time (Month)

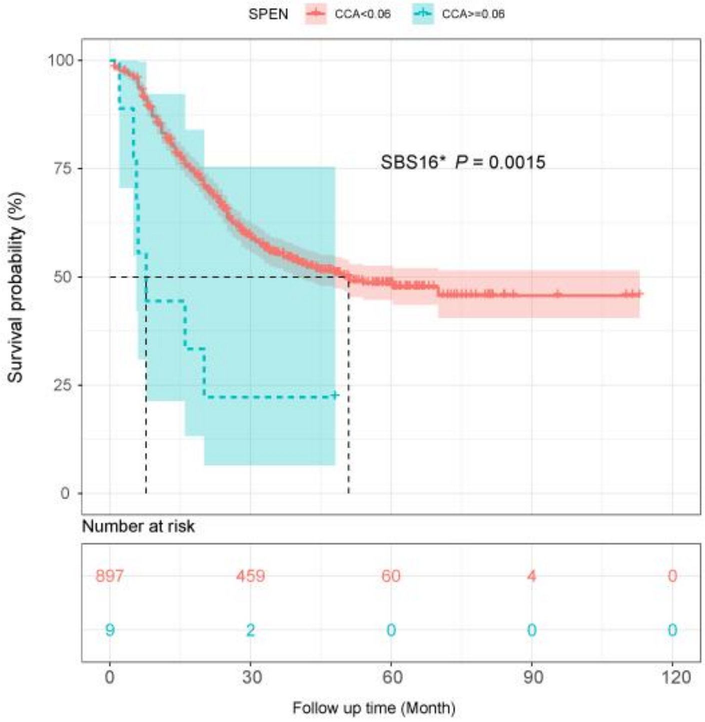
SPEN
CCA<0.06
CCA>=0.06
SBS16* P = 0.0015
Survival probability (%)
100
75
50
25
0
Number at risk
897 459 60 4 0
9 2 0 0 0
0 30 60 90 120
Follow up time (Month)

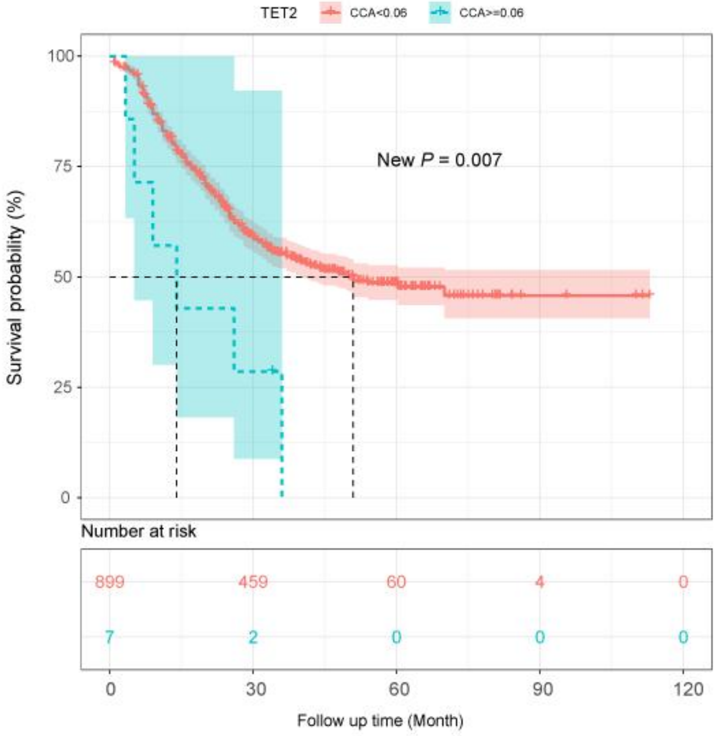
TET2
CCA<0.06
CCA>=0.06
New P = 0.007
Survival probability (%)
100
75
50
25
0
Number at risk
899 459 60 4 0
7 2 0 0 0
0 30 60 90 120
Follow up time (Month)

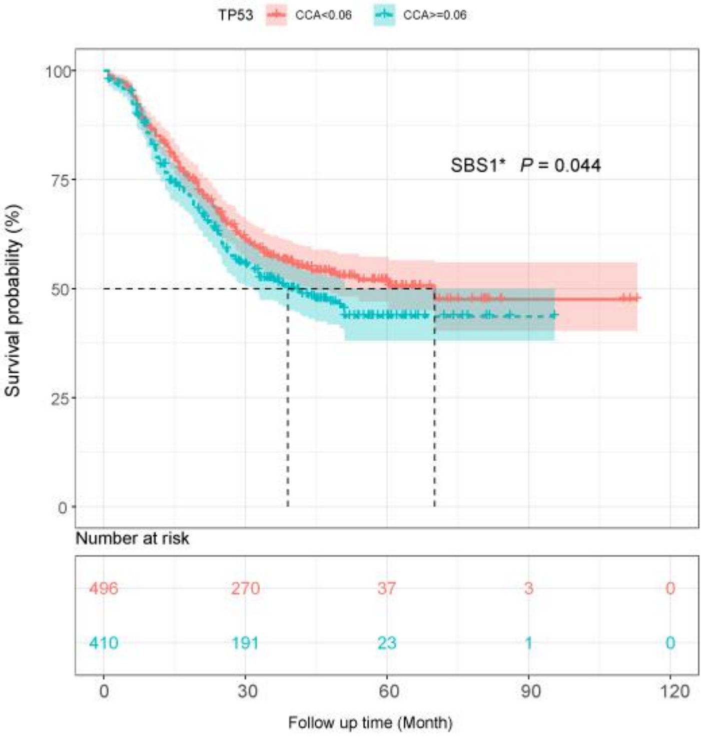
TP53
CCA<0.06
CCA>=0.06
SBS1* P = 0.044
Survival probability (%)
100
75
50
25
0
Number at risk
496 270 37 3 0
410 191 23 1 0
0 30 60 90 120
Follow up time (Month)

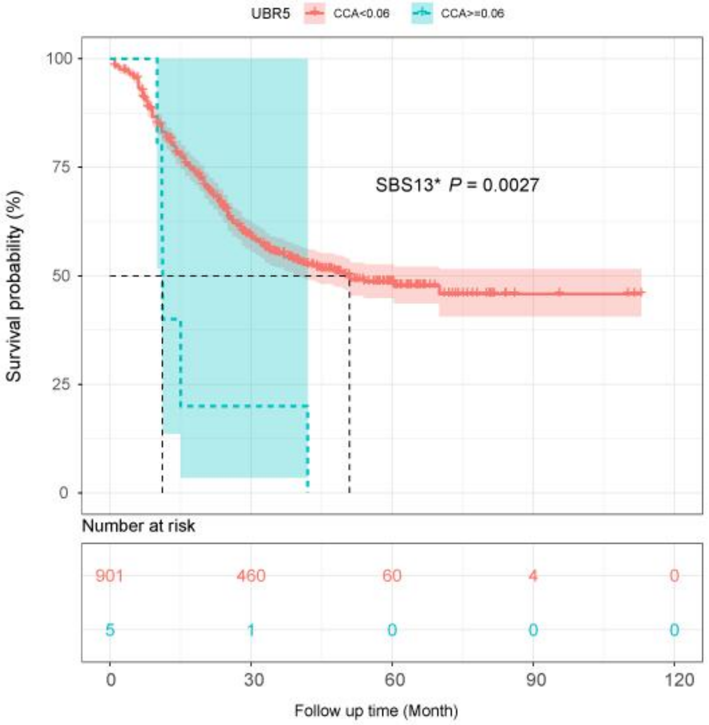
UBR5
CCA<0.06
CCA>=0.06
SBS13* P = 0.0027
Survival probability (%)
100
75
50
25
0
Number at risk
901 460 60 4 0
5 1 0 0 0
0 30 60 90 120
Follow up time (Month)

b

Multivariate Cox Analysis

### SBS1* Signature

| Variable | HR (95% CI) | P−value |
|---|---|---|
| Age | 1.01 (0.997 ~ 1.03) | 0.119 |
| Gender(Male vs. Female) | 1.08 (0.85 ~ 1.38) | 0.517 |
| Stage (I,II,III,IV) | 2.37 (1.91 ~ 2.95) | 5.23e−15 |
| CASP8-SBS1*(CCA : >=0.06 vs. <0.06) | 3.1 (1.36 ~ 7.07) | 0.00727 |
| PTPRC-SBS1*(CCA : >=0.06 vs. <0.06) | 2.25 (0.662 ~ 7.65) | 0.194 |
| RNF213-SBS1*(CCA : >=0.06 vs. <0.06) | 3.86 (1.91 ~ 7.82) | 0.000177 |
| KAT6B-SBS1*(CCA : >=0.06 vs. <0.06) | 1.46 (0.497 ~ 4.31) | 0.49 |
| CIC-SBS1*(CCA : >=0.06 vs. <0.06) | 3.08 (1.25 ~ 7.56) | 0.0144 |
| NFE2L2-SBS1*(CCA : >=0.06 vs. <0.06) | 3.53 (2.01 ~ 6.18) | 1.05e−05 |
| TP53-SBS1*(CCA : >=0.06 vs. <0.06) | 1.01 (0.818 ~ 1.25) | 0.915 |

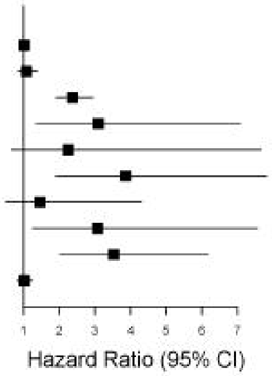


### SBS2* Signature

| Variable | HR (95% CI) | P−value |
|---|---|---|
| Age | 1.01 (0.999 ~ 1.03) | 0.0719 |
| Gender(Male vs. Female) | 1.07 (0.841 ~ 1.36) | 0.579 |
| Stage (I,II,III,IV) | 2.39 (1.93 ~ 2.97) | 2.54e−15 |
| CBL-SBS2*(CCA : >=0.06 vs. <0.06) | 3.08 (1.14 ~ 8.31) | 0.0264 |
| PIK3CA-SBS2*(CCA : >=0.06 vs. <0.06) | 0.958 (0.608 ~ 1.51) | 0.852 |

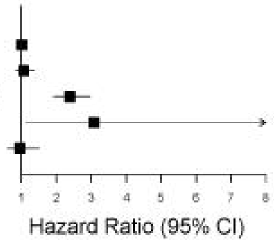


### SBS3* Signature

| Variable | HR (95% CI) | P−value |
|---|---|---|
| Age | 1.01 (0.998 ~ 1.03) | 0.098 |
| Gender(Male vs. Female) | 1.06 (0.832 ~ 1.35) | 0.642 |
| Stage (I,II,III,IV) | 2.39 (1.93 ~ 2.97) | 2.09e−15 |
| CDH10-SBS3*(CCA : >=0.06 vs. <0.06) | 4.16 (1.96 ~ 8.87) | 0.000217 |

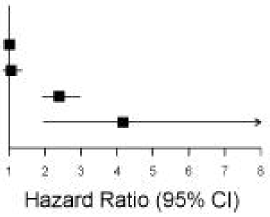


### New Signature

| Variable | HR (95% CI) | P−value |
|---|---|---|
| Age | 1.01 (0.998 ~ 1.03) | 0.092 |
| Gender(Male vs. Female) | 1.09 (0.853 ~ 1.39) | 0.5 |
| Stage (I,II,III,IV) | 2.42 (1.95 ~ 3.01) | 9.67e−16 |
| SNX29-New(CCA : >=0.06 vs. <0.06) | 5.16 (1.63 ~ 16.4) | 0.00529 |
| CHD2-New(CCA : >=0.06 vs. <0.06) | 5.18 (1.64 ~ 16.3) | 0.00502 |
| TET2-New(CCA : >=0.06 vs. <0.06) | 1.46 (0.544 ~ 3.94) | 0.45 |

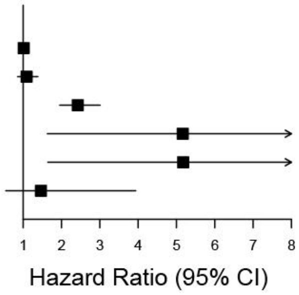

### SBS5* Signature

| Variable | HR (95% CI) | P-value |
|---|---|---|
| Age | 1.01 (0.998 ~ 1.03) | 0.0909 |
| Gender(Male vs. Female) | 1.07 (0.844 ~ 1.37) | 0.561 |
| Stage (I,II,III,IV) | 2.4 (1.94 ~ 2.98) | 1.49e−15 |
| PRDM1-SBS5*(CCA : >=0.06 vs. <0.06) | 3.92 (1.74 ~ 8.85) | 0.000984 |
| RNF213-SBS5*(CCA : >=0.06 vs. <0.06) | 4.68 (2.31 ~ 9.5) | 1.87e−05 |

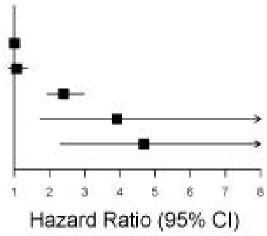


### SBS13* Signature

| Variable | HR (95% CI) | P-value |
|---|---|---|
| Age | 1.01 (0.996 ~ 1.02) | 0.178 |
| Gender(Male vs. Female) | 1.06 (0.835 ~ 1.36) | 0.613 |
| Stage (I,II,III,IV) | 2.46 (1.98 ~ 3.06) | 4.79e−16 |
| UBR5-SBS13*(CCA : >=0.06 vs. <0.06) | 2.57 (0.951 ~ 6.92) | 0.0627 |
| ARHGAP5-SBS13*(CCA : >=0.06 vs. <0.06) | 2.33 (1.03 ~ 5.27) | 0.042 |
| BRCA1-SBS13*(CCA : >=0.06 vs. <0.06) | 3.13 (1.39 ~ 7.05) | 0.00603 |
| SETD2-SBS13*(CCA : >=0.06 vs. <0.06) | 5.67 (1.77 ~ 18.2) | 0.00345 |
| PIK3CA-SBS13*(CCA : >=0.06 vs. <0.06) | 0.806 (0.479 ~ 1.35) | 0.415 |

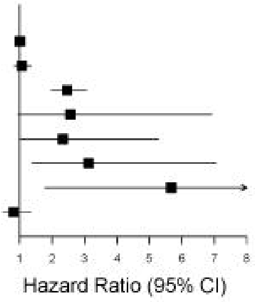


### SBS15* Signature

| Variable | HR (95% CI) | P-value |
|---|---|---|
| Age | 1.01 (0.998 ~ 1.03) | 0.108 |
| Gender(Male vs. Female) | 1.06 (0.833 ~ 1.35) | 0.635 |
| Stage (I,II,III,IV) | 2.42 (1.95 ~ 3) | 8.77e−16 |
| PRDM1-SBS15*(CCA : >=0.06 vs. <0.06) | 4.14 (1.69 ~ 10.1) | 0.00182 |

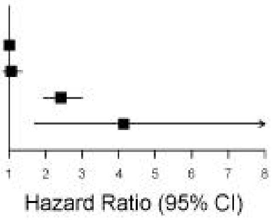


### SBS16* Signature

| Variable | HR (95% CI) | P-value |
|---|---|---|
| Age | 1.01 (1 ~ 1.03) | 0.0536 |
| Gender(Male vs. Female) | 1.06 (0.836 ~ 1.35) | 0.614 |
| Stage (I,II,III,IV) | 2.39 (1.93 ~ 2.97) | 2.3e−15 |
| CDKN2A-SBS16*(CCA : >=0.06 vs. <0.06) | 4.63 (1.68 ~ 12.7) | 0.00299 |
| SPEN-SBS16*(CCA : >=0.06 vs. <0.06) | 2.35 (0.961 ~ 5.76) | 0.0611 |
| NFE2L2-SBS16*(CCA : >=0.06 vs. <0.06) | 3.18 (1.82 ~ 5.56) | 5.08e−05 |

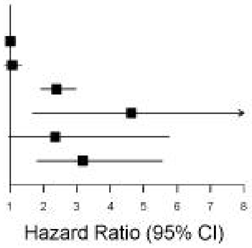


### SBS18* Signature

| Variable | HR (95% CI) | P-value |
|---|---|---|
| Age | 1.01 (0.998 ~ 1.03) | 0.101 |
| Gender(Male vs. Female) | 1.07 (0.843 ~ 1.36) | 0.571 |
| Stage (I,II,III,IV) | 2.38 (1.92 ~ 2.95) | 3.51e−15 |
| PRDM1-SBS18*(CCA : >=0.06 vs. <0.06) | 3.42 (1.35 ~ 8.67) | 0.00942 |
| CASP8-SBS18*(CCA : >=0.06 vs. <0.06) | 2.1 (0.834 ~ 5.26) | 0.115 |
| SETD2-SBS18*(CCA : >=0.06 vs. <0.06) | 2.39 (0.763 ~ 7.45) | 0.135 |

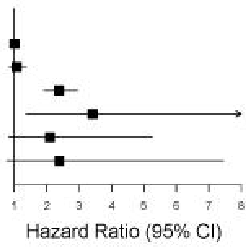

### SBS22* Signature

| Variable | HR (95% CI) | P−value |
|---|---|---|
| Age | 1.01 (0.998 ~ 1.03) | 0.0995 |
| Gender(Male vs. Female) | 1.1 (0.862 ~ 1.4) | 0.452 |
| Stage (I,II,III,IV) | 2.43 (1.96 ~ 3.01) | 5.89e−16 |
| NOTCH1-SBS22*(CCA : >=0.06 vs. <0.06) | 4.28 (1.9 ~ 9.67) | 0.000469 |

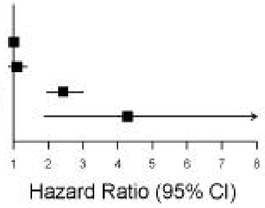


### SBS33* Signature

| Variable | HR (95% CI) | P−value |
|---|---|---|
| Age | 1.01 (0.998 ~ 1.03) | 0.0888 |
| Gender(Male vs. Female) | 1.07 (0.843 ~ 1.37) | 0.563 |
| Stage (I,II,III,IV) | 2.36 (1.9 ~ 2.92) | 7.95e−15 |
| PRDM1-SBS33*(CCA : >=0.06 vs. <0.06) | 4.67 (1.91 ~ 11.4) | 0.000746 |
| RNF213-SBS33*(CCA : >=0.06 vs. <0.06) | 4.84 (1.53 ~ 15.3) | 0.00725 |
| ALK-SBS33*(CCA : >=0.06 vs. <0.06) | 4.12 (1.53 ~ 11.1) | 0.00518 |
| BIRC6-SBS33*(CCA : >=0.06 vs. <0.06) | 3 (1.11 ~ 8.1) | 0.0305 |

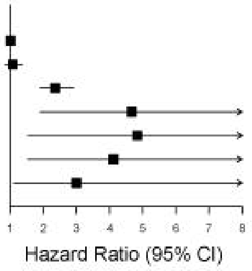
