## Supplementary figures and images for "A practical framework RNMF for the potential mechanism of cancer progression with the analysis of genes cumulative contribution abundance"

### Sup Fig. 2 .pdf

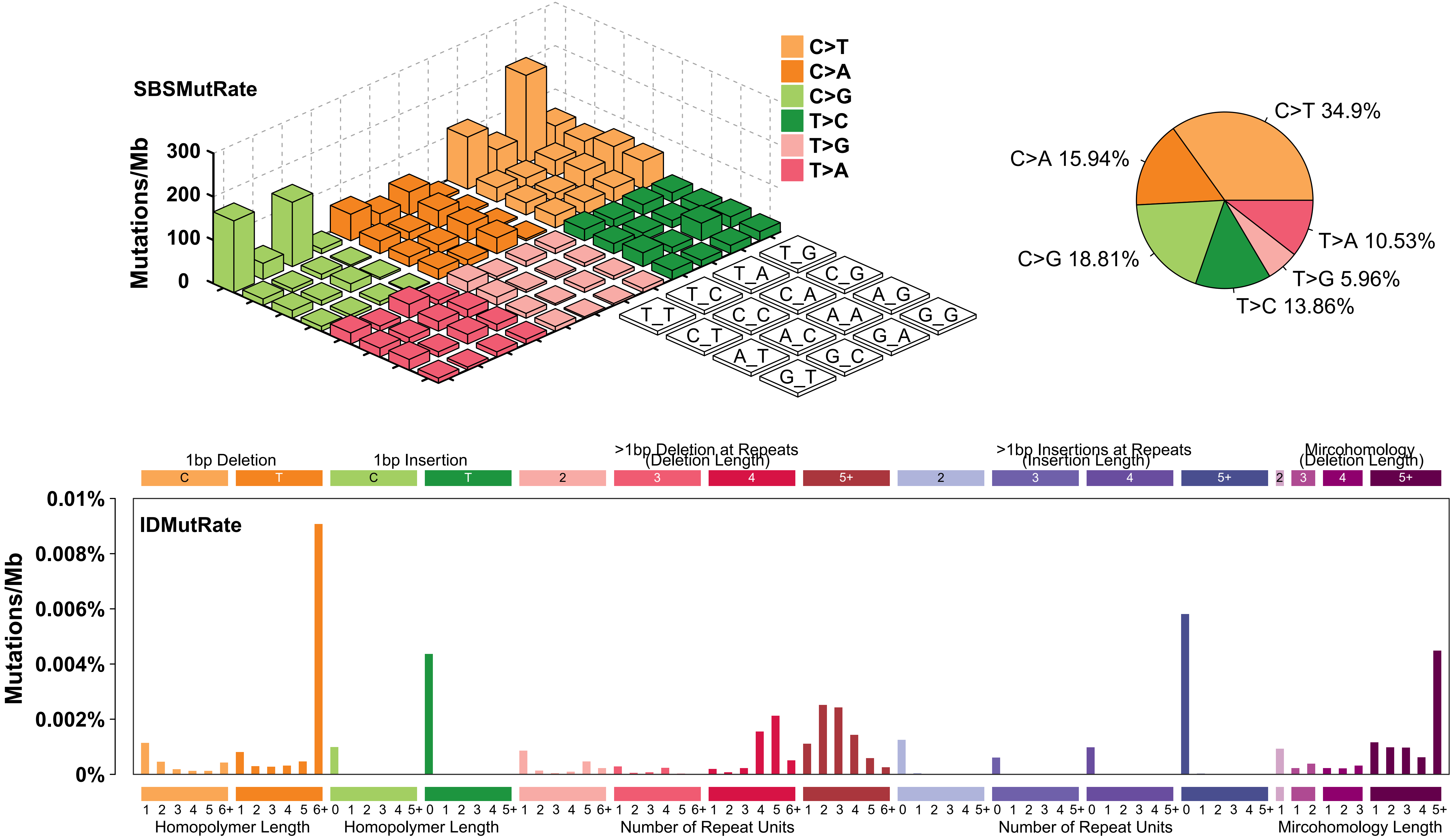

### Sup Fig. 3 .pdf

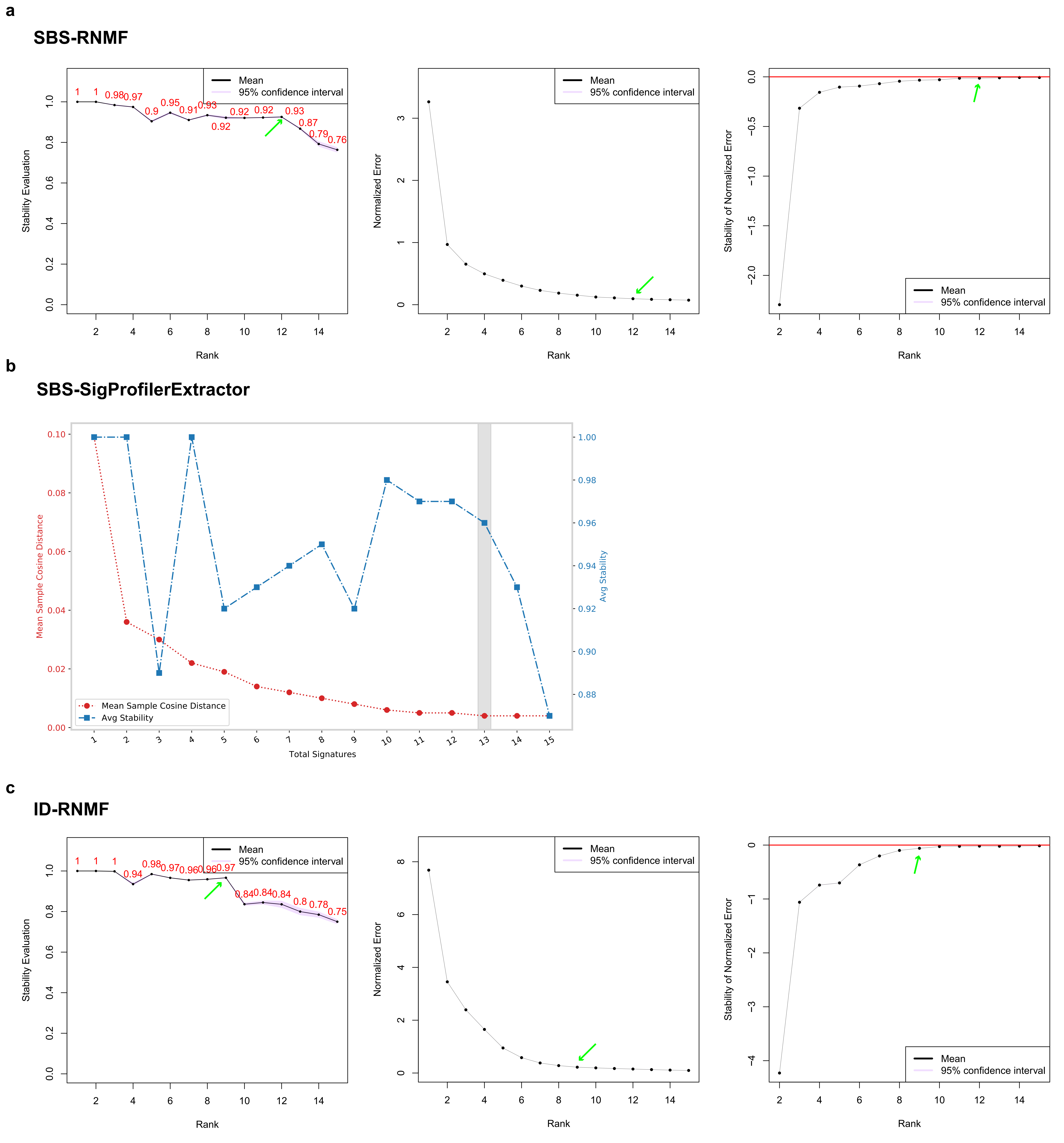

### Sup Fig. 4 .pdf

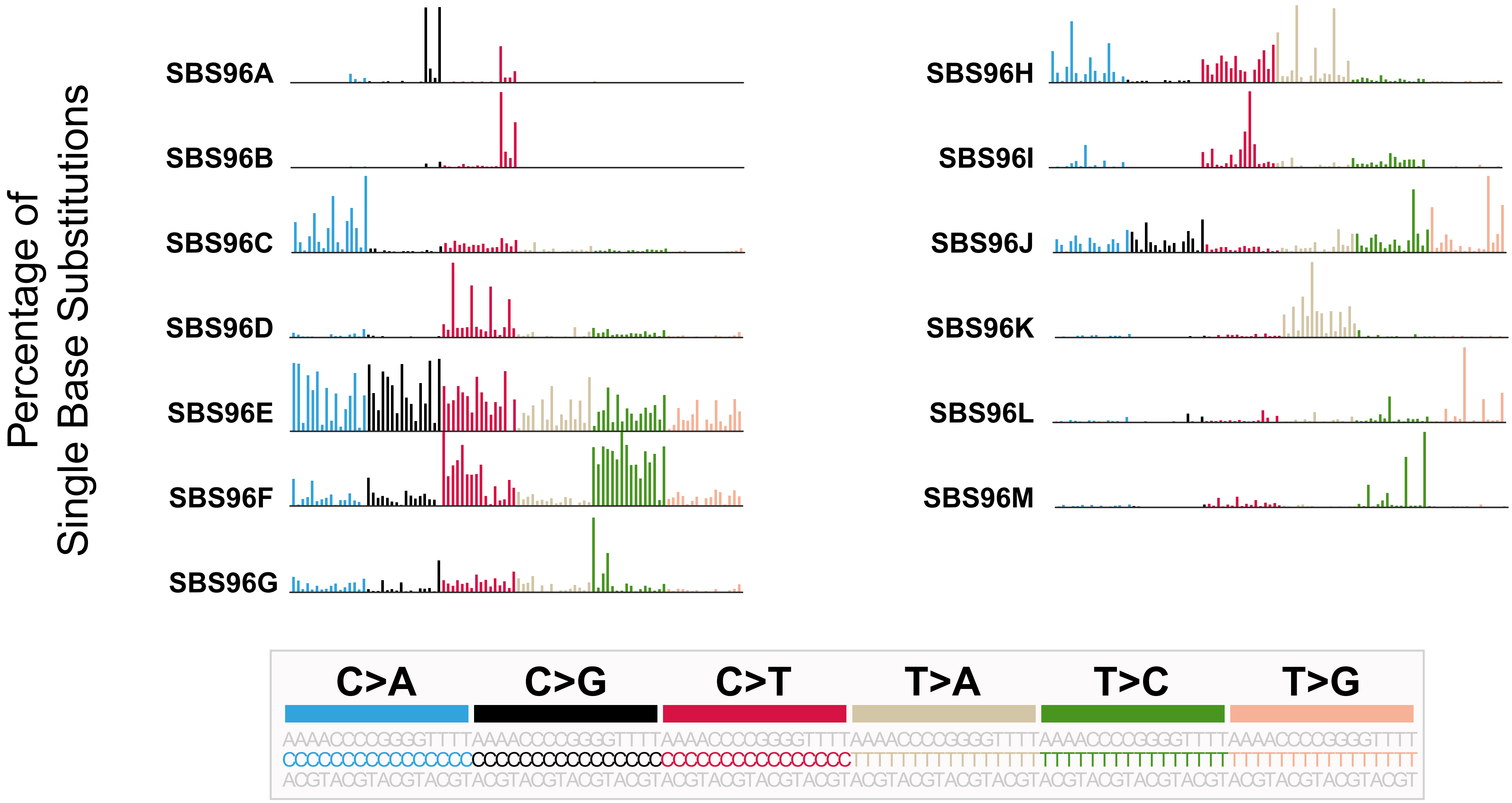

### Sup Fig. 5 .pdf

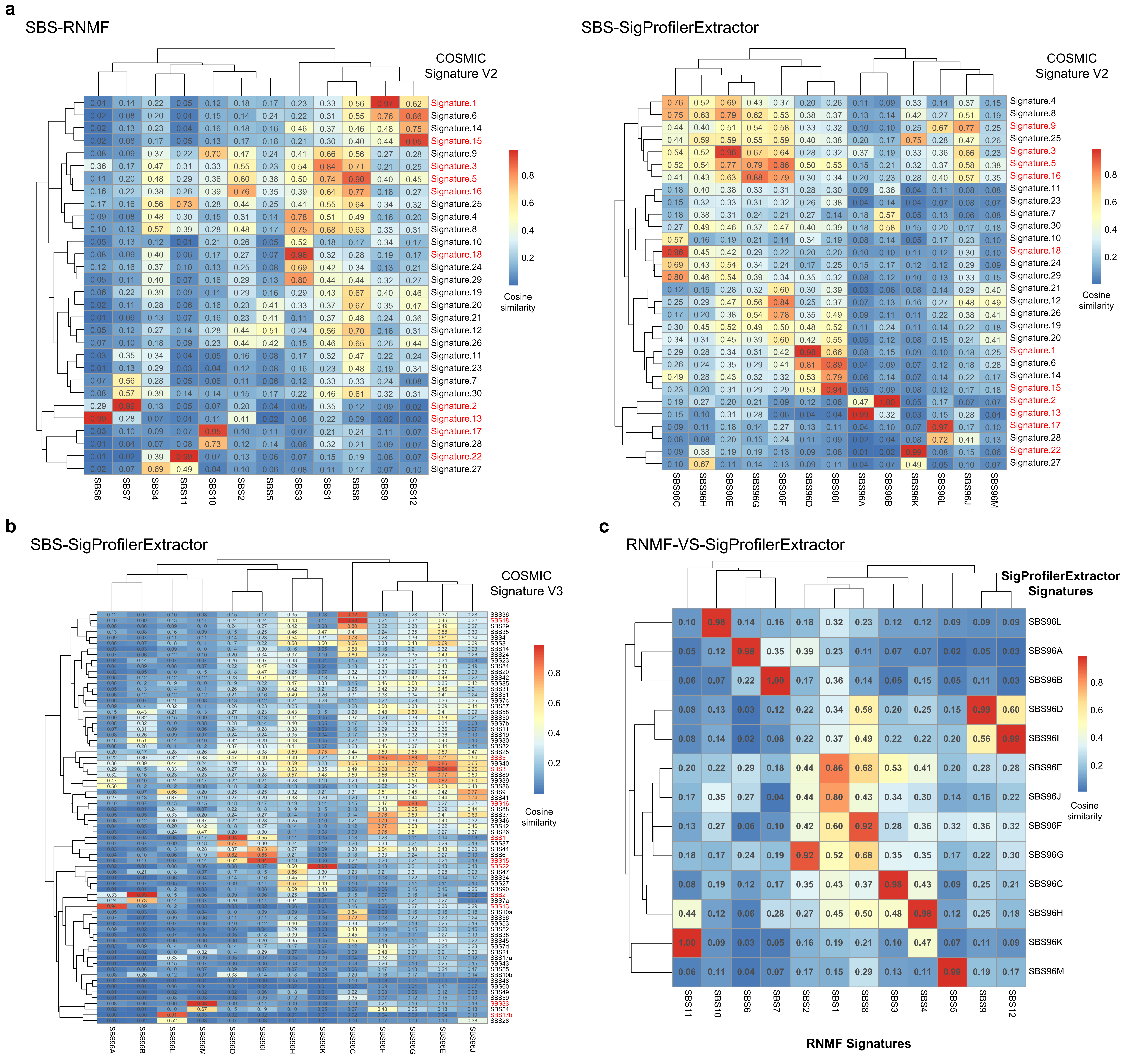

### Sup Fig. 6 .pdf

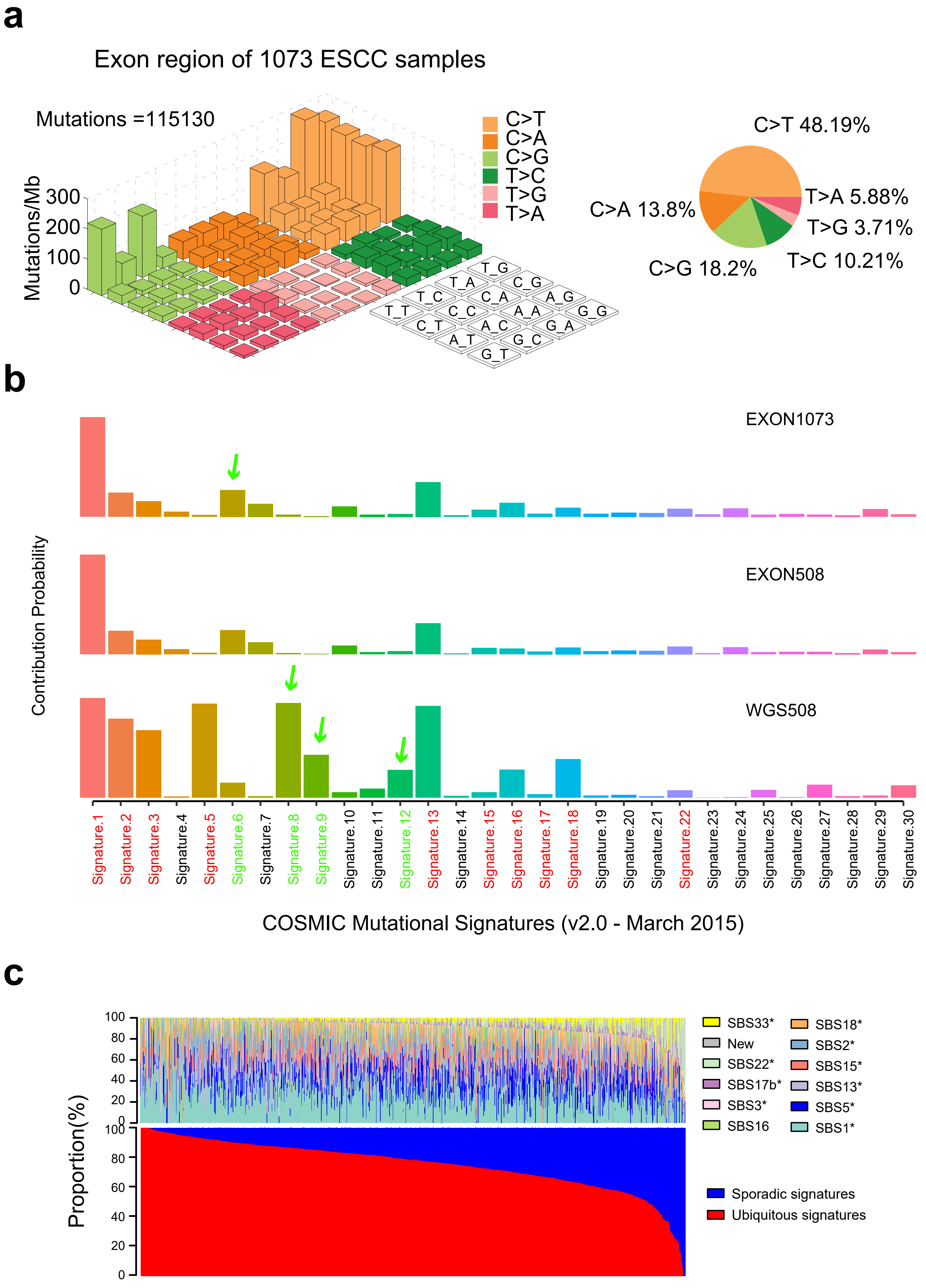

### Sup Fig. 8 .pdf

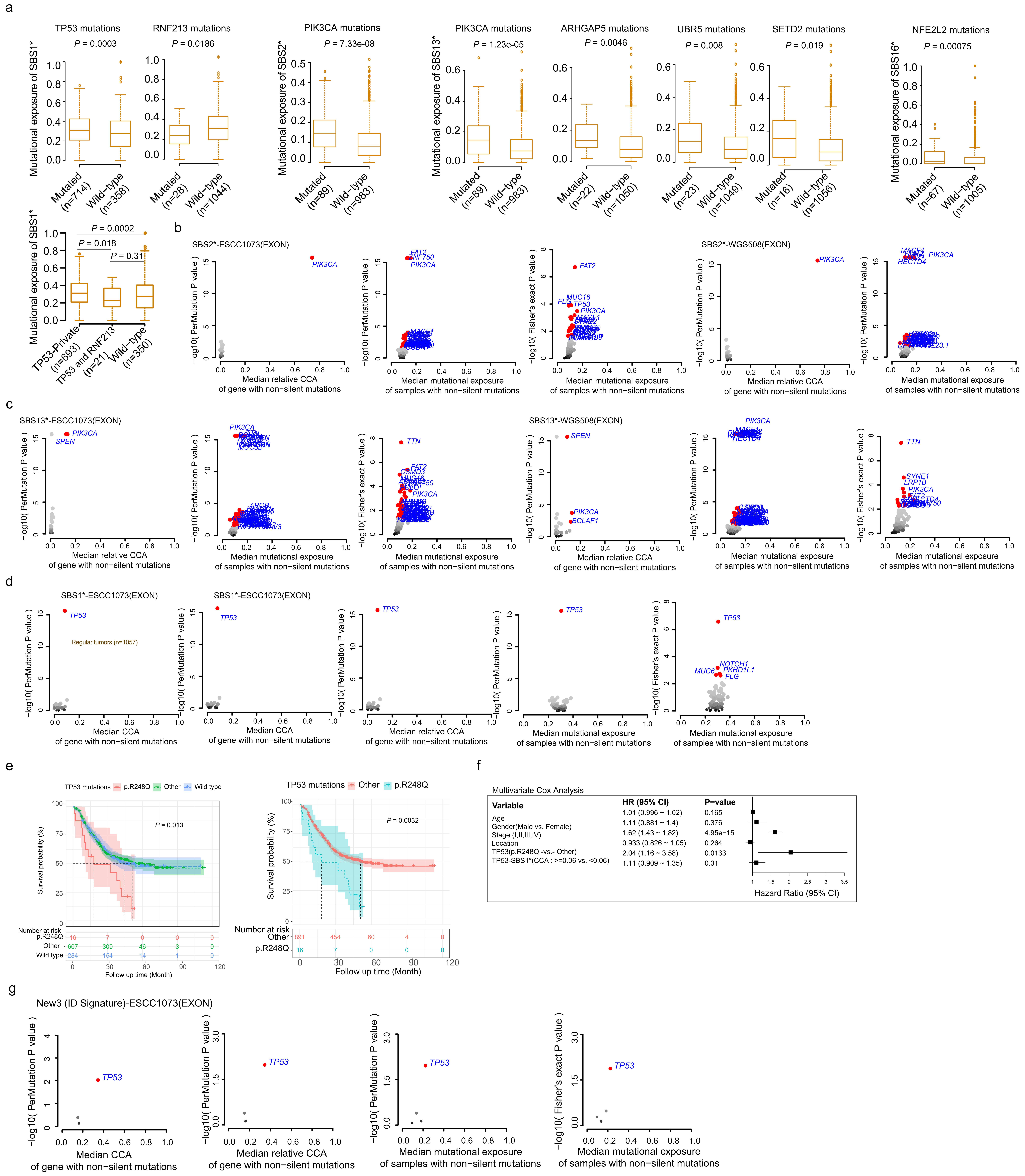

### Sup Fig. 9 .pdf

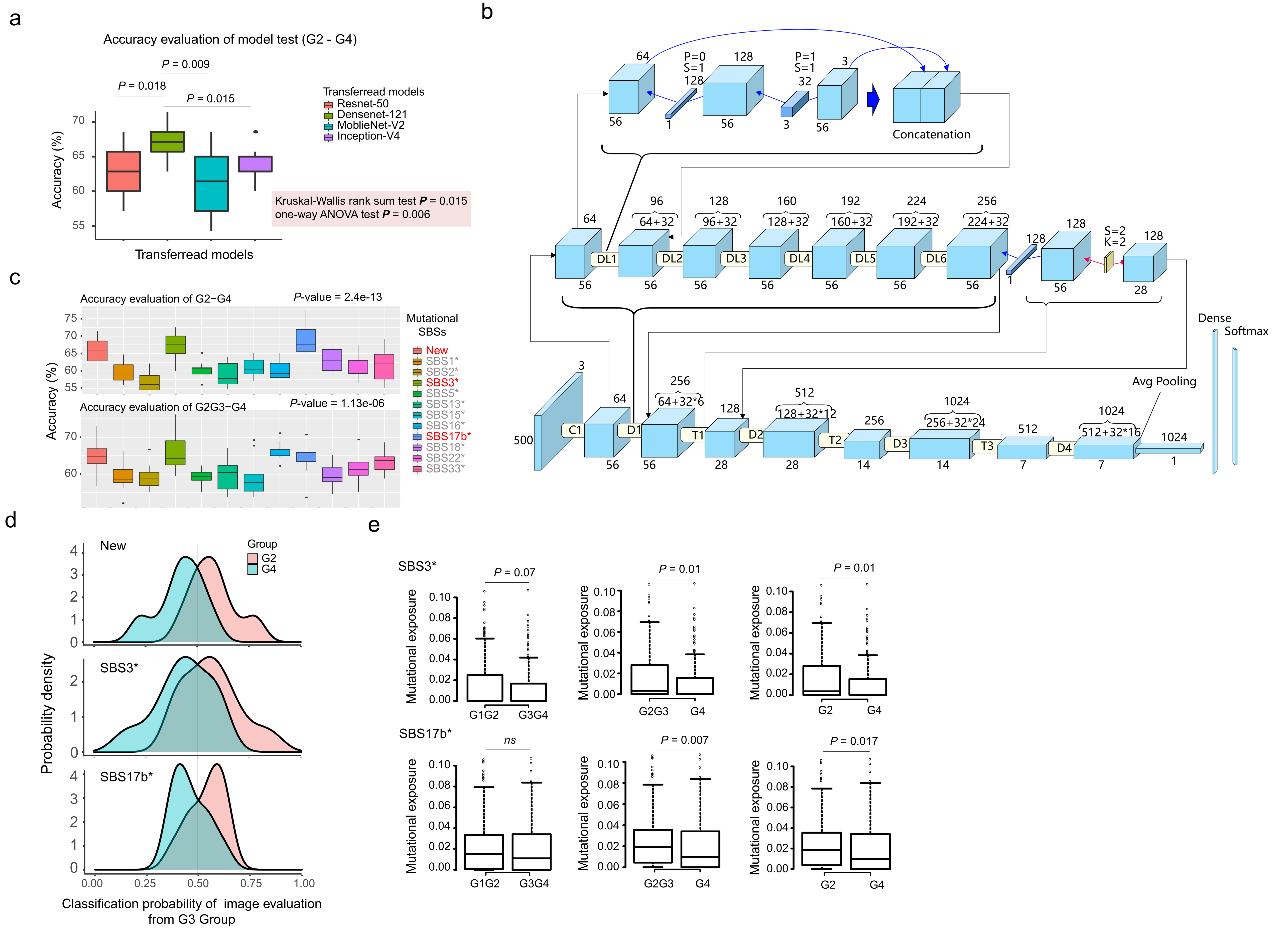
